# A Unified Control of Cellular Differentiation: From Temporal Multistability to Spatial Pattern Formation in Gene Regulatory Networks

**DOI:** 10.64898/2026.04.04.699778

**Authors:** Tejas Bansod, Amrit Kaur, Mohit Kumar Jolly, Ushasi Roy

## Abstract

How genetically identical cells spontaneously break symmetry to assume divergent fates is a fundamental problem in developmental biology. While modern genomics has mapped the vast molecular repertoire involved in gene regulation, understanding the mechanism of cell state transitions that drive differentiation remains a formidable challenge. To address this, we use a reaction-kinetic framework to analyze recurring motifs of two and three competing master regulators. While typically such circuits are studied numerically, we show that assuming symmetry in nodes and interactions provides exact analytical description of the bifurcations governing cell fate transitions. We find that the possible cell fates across all considered topologies are dictated by a single dimensionless quantity, *β*—the ratio of protein degradation to production rates. In the binary Toggle Switch (TS), decreasing *β* destabilizes the symmetric (stem cell) state, giving rise to two asymmetric (differentiated) fates via a supercritical pitchfork bifurcation. In the three-component Toggle Triad (TT), low values of *β* yield three asymmetric fates through subcritical pitchfork bifurcation, creating an intermediate range of *β* where both symmetric and asymmetric fates are simultaneously stable. For the Self-Activating Toggle Switch (SATS), we identify a new parameter for the self-activation threshold (*θ*) and show that decreasing *θ* progressively stabilizes the uncommitted state, leading to a regime of tristability. Building on these temporal bifurcations, we next address the feasibility of spatial structure formation: can these multistable fates stably coexist within a spatial domain? Through a minimal model of cell-cell communication via free diffusion, we extend these motifs into reaction-diffusion systems, which reveals a direct role of network topology on spatial organization. We prove that any heterogeneous pattern in two-node circuits is inherently transient and unstable. In contrast, the three-node repressive network supports the stable spatial coexistence of differentiated phenotypes through pure diffusion, a phenomenon we analyze by studying heteroclinic interface solutions as building blocks. By reducing complex regulatory dynamics to tractable models with physically meaningful parameters, we establish a minimal framework which relates topology to cell fate. Finally, the effects of temporal multistability on pattern formation provide an excellent studying ground for morphogenesis, synthetic biology, and the overarching problem of spatiotemporal self-organization.

## 1. Introduction

Every complex organism begins as a single cell and develops towards its final form through four key processes: **growth, differentiation, pattern formation, and morphogenesis** [1]. Recent advances in omics technologies [2] and computational power [3] have provided unprecedented insight into the chemical basis of these processes, allowing us to map distinct cellular phenotypes to the unique regulation of a common genetic material. At its core, this regulation consists of the selective orchestration of switching genes on or off, which constitutes the chemical basis of differentiation.

But given the scale and complexity of cellular processes and the diversity of functions that must be performed, it is useful to conceptualize these biochemical circuits as information-processing systems divided into sensory inputs, core processing, and phenotypic outputs [4, 5]. At the heart of the core processing layer within gene regulatory networks are recurring **network motifs**—patterns that occur more frequently than expected by chance [6, 7]—which often drive critical cell-fate decisions [8, 9]. In such motifs, a small set of key molecules known as master regulators typically interact through mutual inhibition, creating a competition in which the dominant regulator drives the cell toward a distinct phenotype [10, 11]. Through varying combinations of activating and repressing links, more diverse motifs emerge, each with unique dynamical behaviors and functions, including feedforward loops, autoregulatory circuits, memory switches, and oscillators [4, 12, 13, 14]. Insights into how combinations of simpler motifs are embedded in larger networks have also guided the rational design of synthetic genetic circuits [15].

These decision-making motifs are commonly modeled as systems of coupled chemical reactions [16, 17]. This involves identifying suitable components and kinetic parameters, which is notoriously challenging since *in vivo* reaction rates are rarely measurable [18, 19]. Ensemble-based methods such as RACIPE (RAndom CIrcuit PErturbation) [20] address this by sampling broad parameter ranges and initial conditions to capture possible steady states and phenotypic behaviors. For larger networks where detailed kinetics are infeasible, spin-based Boolean frameworks approximate genes as binary variables governed by logical interaction rules [21, 22], enabling scalable exploration of expression states.

However, a detailed and mechanistic mathematical understanding of these transitions often remains elusive because most biochemical models are inherently “sloppy” [23, 24]. In such systems, the emergent phenotypic landscape is highly sensitive to only a few parameter combinations, while the vast majority of kinetic parameters can fluctuate freely without altering the qualitative dynamics. While computational methods allow for scalable exploration of whole parameter spaces, they do not always identify the specific parameters that actually govern state transitions [25].

To isolate such parameters and gain clearer insights, we propose a new framework: assuming strict symmetry between network nodes. This idealization serves four critical purposes: (1) It significantly reduces the number of parameters needed to describe the system. (2) It puts competing phenotypic outcomes on equal footing, treating them as mere cyclic permutations of each other. (3) The stem cell state can be defined simply as the symmetric solution of the chemical kinetics, which makes many analytical calculations possible. (4) After further simplifications, it allows us to identify the crucial control parameters and map all possible phases and bifurcations in closed form.

Analogous to how the simplicity of the Ising model [26] reveals universal features of phase transitions [27] despite ignoring microscopic details, our symmetric framework provides exact bifurcation conditions. This provides earlier numerical observations with more rigorous foundations and emphasizes the fundamental role of symmetry in uncovering the governing principles of complex dynamical systems [28, 29].

In developing this framework, we consider three archetypal regulatory motifs: the binary toggle switch (TS) of two mutually inhibiting genes; the toggle triad (TT) of three mutually inhibiting genes; and the self-activating toggle switch (SATS), where mutual inhibition is combined with self-activation. We select these specific motifs because they have been extensively studied using numerical techniques [16, 30, 31], providing a clear benchmark for our analytical results.

We note, however, that this reaction-kinetic modeling addresses the chemical fates of a single isolated cell. In functional tissues, cells must coordinate their gene-expression states across space [32]. This coordination can arise through diffusible signals [33], direct exchange via gap junctions [34], and contact-dependent interactions between neighboring cells [35], generating spatial variation in gene activity and leading to pattern formation [32, 33]. Since the goal of this paper is to employ the most fundamental modeling approaches, we include this signaling in our framework strictly through the direct spatial diffusion of the interacting chemicals.

Building on the spatial “tug-of-war” mechanisms explored by Gomez *et al*. [36] and the kinetics of phase ordering and domain coarsening classically studied by Puri and Bray [37, 38], we embed these reaction-kinetic motifs into a system of coupled nonlinear partial differential equations (PDEs). This rigorous PDE analysis allows us to determine whether the multistability generated by these local motifs can autonomously sustain stable spatial interfaces across a tissue, or if any pattern is forced eventually to undergo complete homogenization. This is a highly relevant question since mutual inhibition is a ubiquitous motif in gene networks [39], and the capacity for stable spatial organization is critical. For example, such mechanisms underlie stripe formation in the *Drosophila* embryo [40], where mutually repressive gap-gene networks establish pre-patterns that guide subsequent morphogenesis [41] and the emergence of distinct body structures [42].

Taken together, we wish to schematize a gene-regulatory framework that connects differentiation and its controllability to pattern formation and, ultimately, to the emergence of biological form. The remainder of this paper is organized as follows. Section 2 develops the symmetric reaction-kinetic models and analyzes their exact steady states and bifurcation structures. Section 3 extends the framework to spatially coupled systems through PDE analysis of the interface formation. Finally, Section 4 discusses the biological implications of these results and concludes with possible extensions and limitations of the present approach.

## 2. Temporal Multistability in Symmetric GRNs

### 2.1. Binary Fates in the Toggle Switch

The most basic cell fate decision is a binary one. In metazoan development, this occurs first during blastocyst formation, where the embryo differentiates and segregates into the outer trophectoderm and the inner cell mass [43]. From a gene regulatory perspective, this bifurcation is governed by the transcription factors *Cdx2* and *Oct3/4* [44], which act as master regulators for distinct cell fates. Since they repress one another in a mutual inhibition motif [44], exclusivity of cell identity is assured, which precludes hybrid states and ensures a clear choice between two alternative lineages [45].

Even beyond this initial embryonic event, such binary decisions repeat at every level of maturation. This successive lineage commitment forms a hierarchical, branching tree of development [45], extending down to terminal fate choices such as the divergence of erythroid and myeloid blood cells in hematopoiesis, which is again driven by a pair of mutually repressive transcription factors, *GATA-1* and *PU*.*1* [46, 47].

The chemical basis of this mutual inhibition is the genetic toggle switch. This architecture can be represented as a minimal reaction-kinetic system where Gene 1 and Gene 2 produce Protein X and Protein Y, respectively (Fig. 1a). In this configuration, each protein functions as a transcriptional repressor that binds to the regulatory control region of its opposing gene, thereby impeding its production. This mutual antagonism (Fig. 1b), a motif famously realized synthetically in *E. coli* [16], functions dynamically as a nonlinear switch. The mutual repression ensures that any increase in one protein leads to a further decrease in the other, destabilizing the uncommited state and triggering a transition into one of two distinct, stable identities (Fig. 1c).

**Figure 1:**
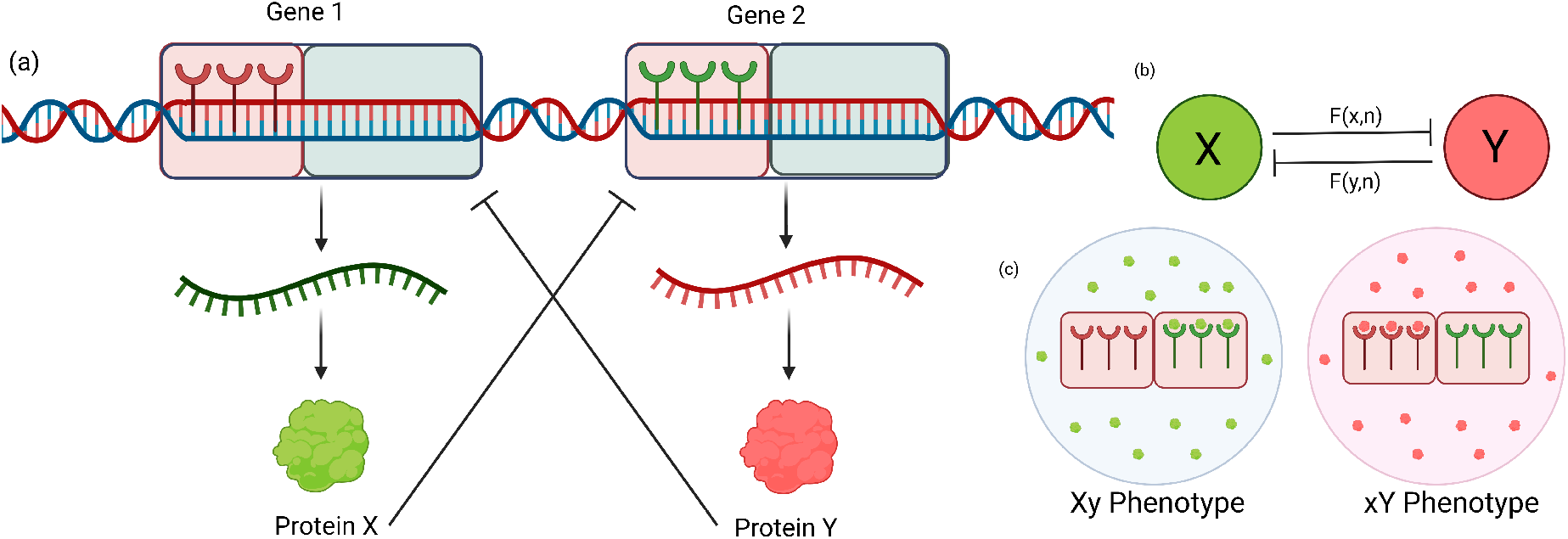
Features of Toggle Switch. (a) Schematic of the gene regulatory circuit where Gene 1 and Gene 2 produce proteins X and Y, respectively. The control region (red) contains binding sites for repression by the opposing protein. The coding (blue) region is transcribed and translated. (b) Network motif representation exhibiting mutual repression: X inhibits production of Y, Y inhibits production of X. (c) Emergence of mutually exclusive phenotypes (*Xy* and *xY* ). One protein completely halts the production of the opposing protein. Construction of synthetic toggle switch in *E. coli* [16] validated this architecture experimentally. Created with Biorender.com [48]

To describe the dynamics of the toggle switch mathematically, we identify two variables, *x* and *y*, representing the protein concentrations of X and Y. These concentrations serve as direct measures of the activation levels for Gene 1 and Gene 2, respectively. While a complete mechanistic model would also track the concentrations of intermediate mRNA molecules, these transcripts typically degrade and reach equilibrium on a much faster timescale than their associated proteins [49]. Consequently, the transcription and translation processes can be subsumed into a single effective production rate via a quasi-steady-state approximation [50]. Following the standard reaction-kinetic framework for biological switches [51, 52], the rate of change for each protein in this two-variable model is determined by a production rate that depends on the concentration of its antagonist, balanced by a linear decay term:

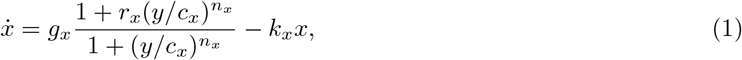

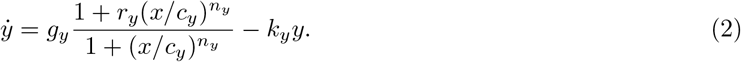

This formulation utilizes Hill-type kinetics to capture the nonlinearities of transcriptional binding [52]. The parameters can be classified into two categories, where (*g, k*) define the reaction speeds of the system; *g* represents the maximal production rate when the gene is fully expressed, and *k* captures the degradation rate due to natural protein breakdown and dilution from cell growth [53]. While the Hill interaction parameters (*c, r, n*) describe the binding interaction between the receptor and repressor ligand, the threshold *c* represents the repressor concentration required to halve the promoter’s maximal activity. The parameter *r* (0 ≤ *r <* 1) accounts for basal or “leaky” transcription [52, 54], reflecting the fact that even fully repressed genes can maintain a residual level of expression. Finally, the Hill coefficient (*n*) dictates the steepness, or sigmoidicity, of the gene’s response. Rather than an exact count of physical binding sites, *n* acts as an upper limit estimate of cooperative binding [52]; for example, *n* = 2 corresponds to dimeric binding, where the attachment of one ligand accelerates the binding of the second, yielding steeper repression.

In its most general formulation, the system described by Equations (1) and (2) contains ten independent parameters. While these are necessary to faithfully model the intricacies of an asymmetric, leaky *in vivo* circuit, their exact values are not easy to measure [25] and fluctuate significantly across cell populations [55]. Fortunately, the capacity of this circuit to execute binary decisions is a robust consequence of its mutual-inhibition topology rather than the precise fine-tuning of individual kinetic rates [56]. In many biological contexts, even the master regulator itself is an abstraction for an entire “team” of co-expressed genes that mutually inhibit opposing teams to enforce a stable phenotype [57]. Because this qualitative behavior is preserved across broad parameter regimes [16], we adopt an abstract view of the toggle switch as a formal dynamical system. This allows us to distill the network down to a simpler form and identify the bifurcation parameters that govern the stability of cell states.

Our first simplification is to treat both genes in Fig. 1 as perfectly symmetric. The toggle switch equations considered by Gardner et al. [16] can be viewed as a specific case of the general formulation (1, 2) where decay rates are unity (*k*_*x*_ = *k*_*y*_ = 1), thresholds are normalized (*c*_*x*_ = *c*_*y*_ = 1), and leakiness is absent (*r* = 0). To further simplify our symmetric model, we make similar assumptions, normalize the threshold (*c* = 1), and assume full repression (*r* = 0). By defining a dimensionless parameter *β* = *k/g*, we arrive at a minimal system governed by the following dimensionless differential equations:

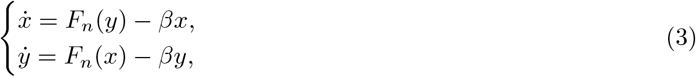

where the nonlinear repression function *F*_*n*_ is defined as:

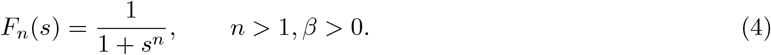

By reducing the system to only two tunable parameters (*β, n*), we show in Proposition 1 (proof in Appendix A) that the model admits up to three steady states: one symmetric and two asymmetric.

##### Proposition 1.

*Consider the toggle switch system defined by* (3) *with the repression function given by* (4). *The system exhibits the following equilibrium, bifurcation, and stability properties:*

1. ***Existence:*** *For every β >* 0, *the system admits a unique symmetric equilibrium* (*ϕ, ϕ*), *where ϕ is the positive real root of the equation βϕ* = *F*_*n*_(*ϕ*).
2. ***Bifurcation:*** *The system undergoes a supercritical pitchfork bifurcation at a critical threshold β*_*c*_, *corresponding to a critical concentration ϕ*_*c*_:

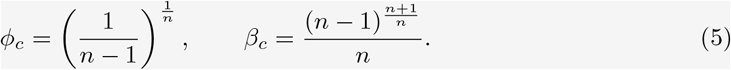
3. ***Stability:*** *For β > β*_*c*_, *the symmetric equilibrium* (*ϕ, ϕ*) *is a global attractor. For β < β*_*c*_, *the symmetric state becomes an unstable saddle point, giving rise to two stable asymmetric equilibria*.

To visualize the dynamical regimes established in Proposition 1, we examine the phase space geometry through nullcline analysis. As illustrated in Figure 2, the system’s steady states are determined by the intersections of the nullclines 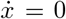 and 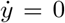. For the representative case *n* = 2 (where *β*_*c*_ = 0.5), the topology of the phase plane shifts qualitatively across the bifurcation threshold. At low degradation rates (*β* = 0.3 in Fig. 2a), the nullclines intersect at three points: a central unstable saddle point and two stable asymmetric nodes, representing the mutually exclusive fates of a differentiated system. And when the relative degradation rate is increased above the threshold (*β* = 0.7 in Fig. 2b), these asymmetric equilibria vanish, leaving only the stable symmetric fixed point that corresponds to the uncommitted progenitor state.

**Figure 2:**
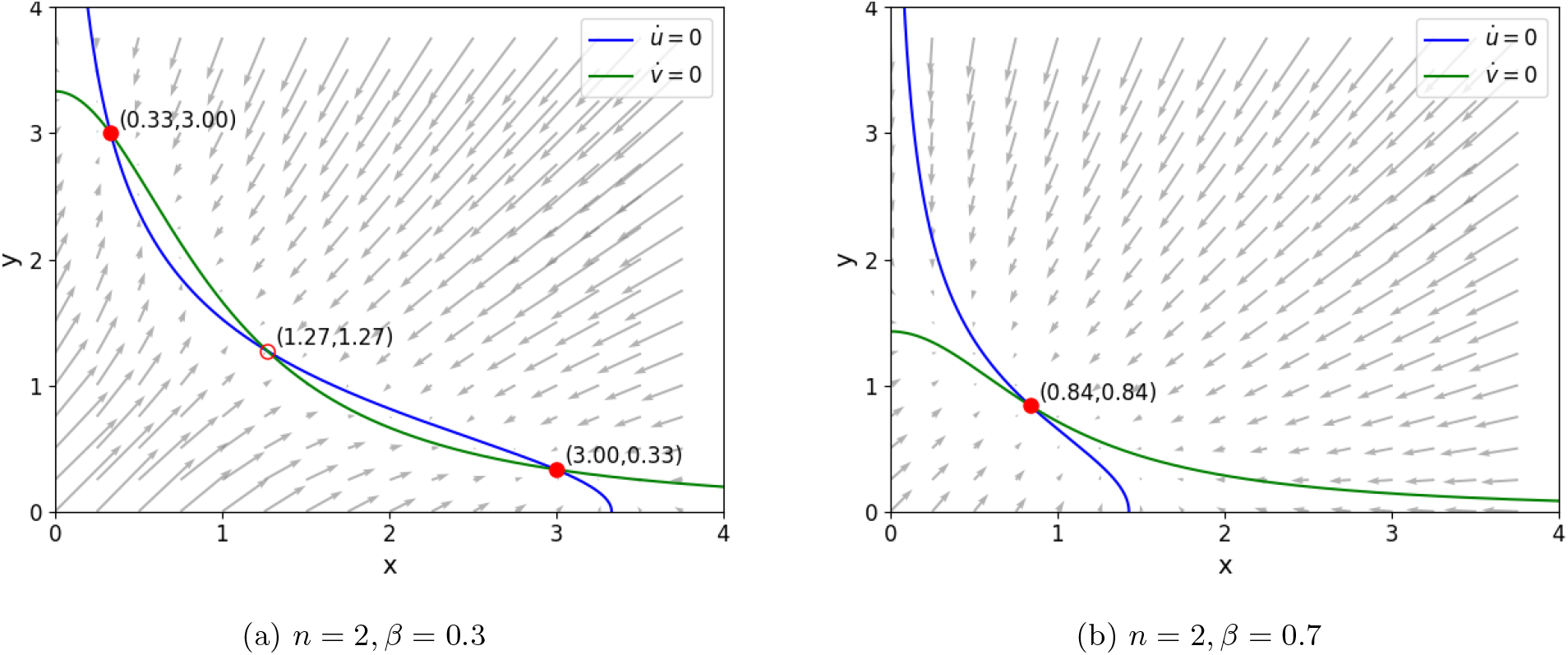
Nullclines and vector fields for Toggle Switch. Plotted for Hill coefficient *n* = 2, with critical parameters *ϕ*_*c*_ = 1 and *β*_*c*_ = 0.5. (a) For *β* = 0.3 *< β*_*c*_, the system exhibits bistability with two asymmetric equilibria. (b) For *β* = 0.7 *> β*_*c*_, only the symmetric equilibrium remains.

While Figure 2 clearly illustrates the phase space topology, it requires fixing both the cooperativity *n* and the degradation rate *β*. To gain a more global view of the dynamics, we can fix *n* and study the continuous role of varying *β* alone. Although bifurcation diagrams are generally plotted for single-variable systems, we can exploit the inherent symmetry of the toggle switch to project the dynamics onto a single axis, plotting the steady-state expression levels and separating the high and low branches. As established in our analytical proposition, the exchange of stabilities occurs exactly at the tipping point (*β*_*c*_, *ϕ*_*c*_), which perfectly matches the numerically determined bifurcation diagrams in Figures 3a and 3b.

**Figure 3:**
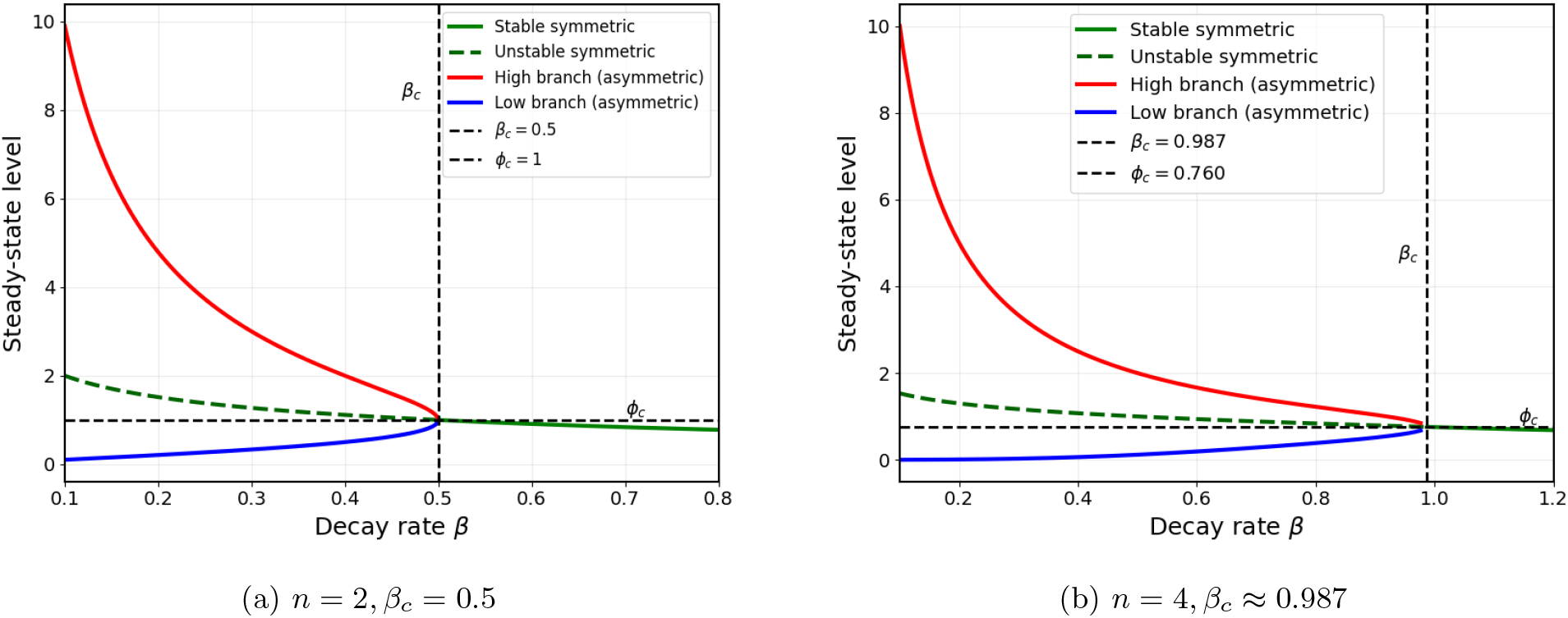
Bifurcation diagrams for the symmetric Toggle Switch. Steady-state expression levels are plotted as a function of the relative degradation rate *β*. The intersection of the dashed lines at (*β*_*c*_, *ϕ*_*c*_) marks the “neck” of the pitchfork, where the symmetric progenitor state (solid green) loses stability (dashed green). Below this critical threshold (*β < β*_*c*_), the system exhibits bistability, diverging into stable high-expression (red) and low-expression (blue) asymmetric branches. As *β* decreases, the widening gap between the two branches signifies an increase in the biochemical distinctness and noise-robustness of the differentiated states. Note however that even for different *n*, the shape of the bifurcation diagrams is structurally the same.

A major strength of the visualization in Fig. 3 is how it highlights the structural basis of the toggle switch: the bifurcation remains a pitchfork across different values of *n*. Furthermore, this offers insight into biological function. Just below the critical threshold *β*_*c*_, the high and low expression branches remain so close that given inherent reaction noise, these small differences would be insufficient to prevent stochastic flips. However, as we reduce *β* further into the bistable regime, the expression gap between the low and high branches widens progressively. A sufficiently large gap ensures that the two asymmetric states represent distinct, committed phenotypes that are stable against the stochasticity of gene expression [58].

Having established how the degradation rate *β* drives the system through a generic pitchfork bifurcation, we now examine how the cooperativity *n* influences this structure. Figure 4 maps the dependence of the critical tipping point (*β*_*c*_, *ϕ*_*c*_) on the Hill coefficient *n*. At the boundary limit of *n* = 1, we find *β*_*c*_ = 0, analytically confirming that direct chemical models require cooperative binding to enforce bistability [51], unless bypassed by alternative mechanisms such as stochastic noise [59] or network degeneracy [60].

**Figure 4:**
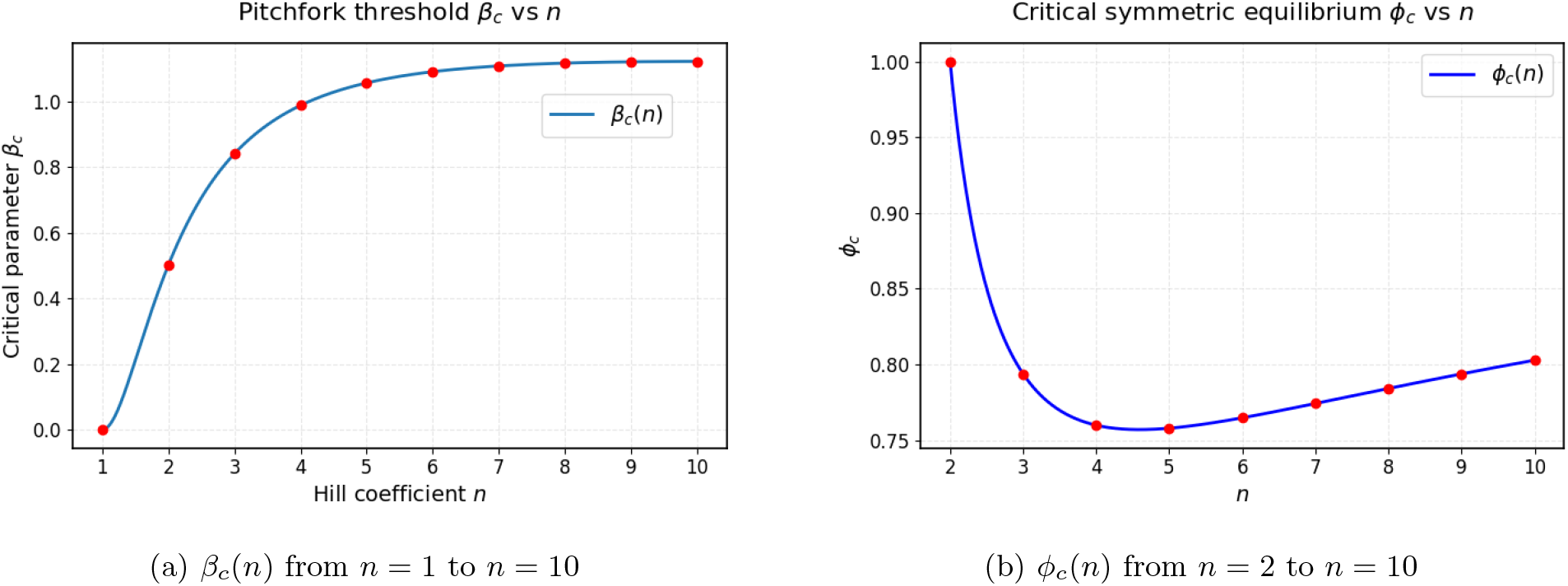
Dependence of critical parameters on cooperativity. (a) The critical threshold *β*_*c*_ increases with the Hill coefficient *n*, with *β*_*c*_ = 0.5 at *n* = 2. This indicates that while higher cooperativity promotes bistability, the effect saturates quickly. (b) The critical symmetric equilibrium *ϕ*_*c*_ exhibits non-monotonic dependence on *n*, reaching a minimum value near *n* ≈ 5. Note that *ϕ*_*c*_ is undefined for *n* = 1, and hence plot starts from *n* = 2.

As *n* increases beyond unity, *β*_*c*_ rises sharply, indicating that higher cooperativity expands the bistable parameter space. This expansion is a well-documented driver of genetic robustness [16, 61], with evolutionary simulations suggesting that such values naturally emerge when selecting for functional states with large stable asymmetries [62]. However, we find that *β*_*c*_(*n*) saturates rapidly beyond *n* ≈ 5, aligning with the fact that natural regulatory motifs typically employ moderate Hill coefficients (*n* ∈ [2, 4]) [63]. While high cooperativity sharpens decision-making, it also amplifies input noise and reduces information processing capacity [64]; thus, moderate cooperativity ensures the functional robustness of the decision-making process by optimizing the trade-off between signal steepness and noise sensitivity.

Finally we note that the critical symmetric equilibrium *ϕ*_*c*_(*n*) displays a qualitatively different, non-monotonic behavior (Fig. 4b). Rather than a simple trend, *ϕ*_*c*_ drops to a global minimum near *n* ≈ 5 before gradually increasing. It remains unclear whether this specific non-monotonicity has any biological significance. However, because maintaining bistable memory across varying protein degradation rates (*β*_*c*_) likely imposes a far stricter evolutionary constraint than the exact stem cell concentration (*ϕ*_*c*_), this behavior may simply be just an artifact rather than of primary importance in toggle switch behavior.

### 2.2. Ternary Fates in the Toggle Triad

While the toggle switch analyzed previously drives simple binary decisions, cells frequently navigate three-way or higher-order branching. They can manage this complexity simply by coupling multiple simpler motifs like toggle switches together. As an example, consider the differentiation of naive CD4^+^ T-helper cells [65]. Upon activation, the progenitor must choose among three distinct phenotypes: regulatory (Treg), pro-inflammatory (Th17), and Th1[66]. This ternary decision is enforced by three corresponding master transcription factors—*Foxp3, RORγt*, and *T-bet* [10]—which mutually repress one another to guarantee a single, exclusive fate. By engaging in this pairwise cross-inhibition, these regulators effectively form a network of interlocked toggle switches known as the toggle triad [31].

Chemically, this architecture is modeled as a reaction-kinetic system where Genes 1, 2, and 3 produce proteins *U, V*, and *W* . Each protein acts as a transcriptional repressor, binding to the control regions of the other two genes to block their production (Fig. 5a). This creates the toggle triad motif, where all three components compete for dominance (Fig. 5b). Dynamically, an initial increase in one protein can drive down the other two, which destabilizes the uncommitted progenitor state. Following the ON/OFF logic of the toggle switch, the system naturally resolves into a state where one gene is fully “ON” and the other two are forced “OFF.” This complete suppression ensures biochemical distinctness, driving a robust transition into one of three mutually exclusive cellular fates (Fig. 5c).

**Figure 5:**
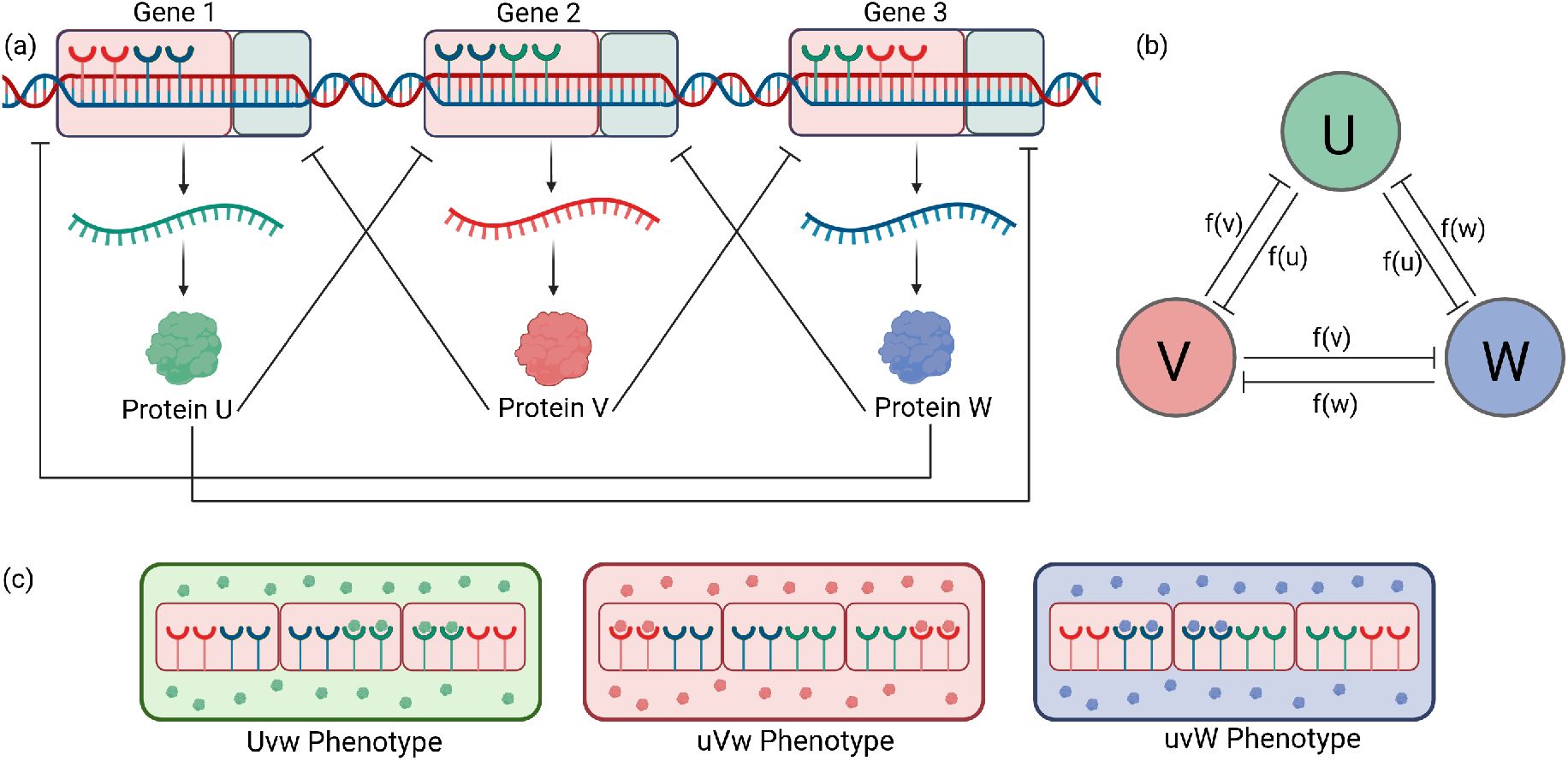
Features of Toggle Triad. (a) Schematic of the gene regulatory circuit where Gene 1, Gene 2, and Gene 3 produce proteins U, V, and W, respectively. The control region (red) contains binding sites for repression by the opposing proteins. The coding (blue) region is transcribed and translated. (b) Network motif representation exhibiting mutual repression: U, V, and W mutually inhibit one another’s production. (c) Emergence of mutually exclusive phenotypes (*Uvw, uV w*, and *uvW* ). Dominance of one protein completely halts the production of the opposing proteins. Created with Biorender.com [48]

Although regulatory networks often exhibit asymmetry—such as uneven interaction strengths or incomplete mutual repression, as seen in the repressive triad of the vertebrate neural tube [67]—we proceed with a perfectly symmetric toggle triad model. To model this mathematically, we extend the dimensionless formulation from the previous section. In our analysis of the binary toggle switch, we observed that the core pitchfork bifurcation persists across different values of the Hill coefficient *n*, tuned entirely by the relative degradation rate *β*. Treating *β* as the primary bifurcation parameter aligns with biological reality: reaction rates (*β*) are dynamically tuned *in vivo*. For example, during CD4^+^ T cell differentiation, external cytokine signals actively increase transcription rates to drive state transitions [10]. Conversely, *n* is a physical property of the ligand-DNA structure and cannot be dynamically tuned. Therefore, as we add the complexity of a third node to our network, we fix *n* and keep *β* as our sole tunable variable.To determine the appropriate value for *n*, we recall that functional switching strictly requires *n >* 1 and natural gene circuits typically operate within a narrow range of *n* between 2 and 4 [63]. From this limited range, we select *n* = 2, which corresponds to the widespread dimeric cooperativity observed in canonical repressors like LacI and TetR [16, 17]. While we expect the overall bifurcation structure to remain independent of the exact value of *n*, the most crucial reason to favor *n* = 2 beyond biological considerations is that the simple quadratic power enables a detailed, closed-form analytical treatment of the system.We define the repression due to dimeric binding as:

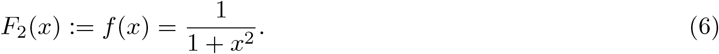

Following standard thermodynamic models of transcription, if the repressive effects from the two competing genes act independently, the probability of promoter occupancy is simply the product of their individual regulation factors [54]. Therefore, the total production rate of a given node is the product of the repression functions of the other two. The dynamics of the toggle triad are then:

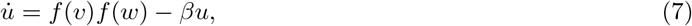

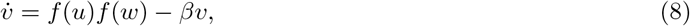

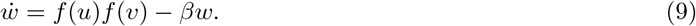

Bifurcation analysis of this model reveals the presence of a symmetric undifferentiated state and three differentiated single positive states. We also consider hybrid states with double positive states, which however turn out to be unfeasible. The stability results for these states are established in Proposition 2 (proof can be found in Appendix B).

##### Proposition 2.

*Consider the symmetric three-node toggle triad defined by Equations (7)–(9), with the dimeric repression function* 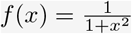 *and relative degradation rate β >* 0. *The system exhibits the following equilibrium and stability properties:*

1. ***Symmetric State (Undifferentiated):***
  - ***Existence:*** *There exists a unique, fully symmetric equilibrium* (*ϕ, ϕ, ϕ*) *satisfying:*

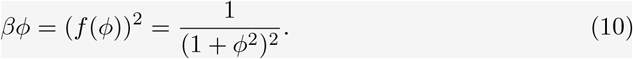
  - ***Stability:*** *This symmetric state is linearly stable if and only if:*

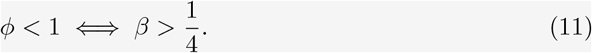
2. ***Asymmetric States (Differentiated):***
  - ***Single Positive States*** (*H, L, L*): *Three distinct asymmetric equilibria—corresponding to permutations of the state* (*H, L, L*) *with one high-expression node* (*H*) *and two low-expression nodes* (*L*)*—exist and are linearly stable if and only if:*

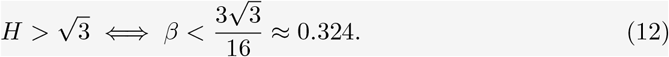
  - ***Hybrid States*** (*H, H, L*): *Three distinct equilibria corresponding to permutations of the state* (*H, H, L*) *with two high-expression nodes and one low-expression node—may exist but are strictly* ***unstable*** *for all valid parameter regimes*.

To see the consequences of Proposition 2, we first plot the bifurcation diagram for the toggle triad. The diagram in Fig. 6 reveals a more complex transition than the simple continuous pitchfork observed in the binary switch. As *β* is decreased from a high value, the symmetric undifferentiated state loses stability at the critical threshold *β*_1_ = 1*/*4. This transition is characteristic of a subcritical pitchfork bifurcation. where, the asymmetric **HLL** branches emerge abruptly. They appear at the higher parameter value *β*_2_ ≈ 0.324 via saddle-node bifurcations and extend backward, creating a discontinuous jump in biochemical expression levels.

**Figure 6:**
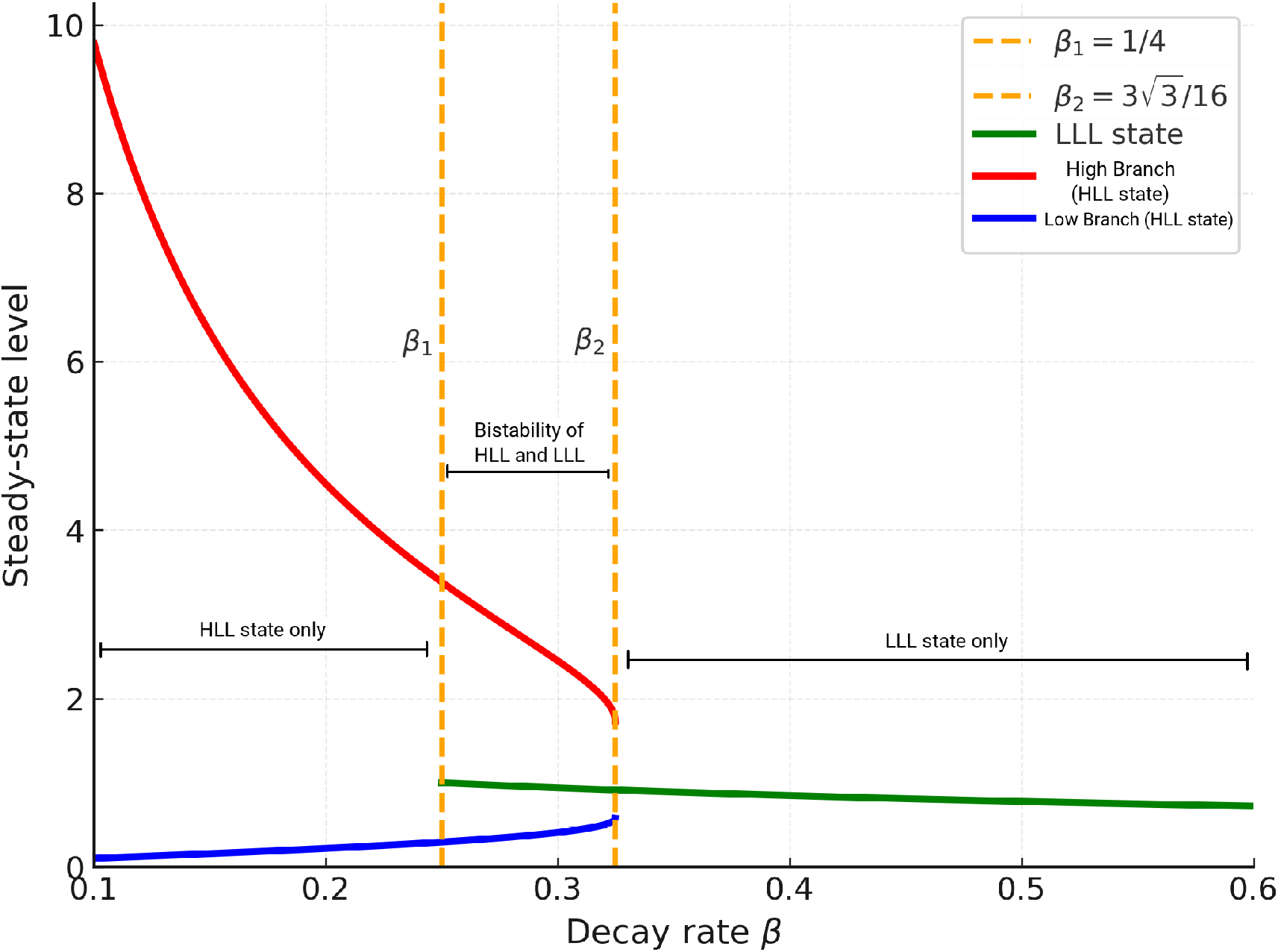
Bifurcation diagram of the symmetric Toggle Triad. Steady-state expression levels are plotted as a function of the relative degradation rate *β*. The solid green line tracks the symmetric, undifferentiated progenitor state (**LLL**), while the solid red and blue lines track the high-expression and low-expression nodes of the differentiated states (**HLL**), respectively. The vertical dashed lines mark the critical stability thresholds at *β*_1_ = 1*/*4 and 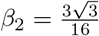. At *β*_1_, the symmetric **LLL** state loses stability (unstable branch extensions are not plotted). For *β < β*_1_, the stable states are all differentiated. For *β > β*_2_ on other extreme, only the uncommitted **LLL** state exists. Crucially, the intermediate window (*β*_1_ ≤ *β* ≤ *β*_2_) represents a regime of multistability, permitting the coexistence of both the **LLL** progenitor and the differentiated **HLL** states.

A critical result of this subcritical bifurcation is the emergence of a coexistence window between *β*_1_ = 0.25 and *β*_2_ ≈ 0.324. In this regime, the system is multistable: the undifferentiated **LLL** progenitor state (green) and the three differentiated **HLL** states (red/blue branches) are simultaneously stable. This phenotypic landscape is summarized in the schematic of Fig. 7, where the light grey progenitor and the colored specialized cells occupy the exact same parameter space. This window provides a mathematical basis for lineage plasticity [10]. Within these bounds, a cell can reside in the progenitor state or, under the influence of molecular noise [58], transition into a committed lineage without requiring a change in the underlying degradation rate. When *β* drops below *β*_1_, the **LLL** state loses its stability, forcing the cell to definitively adopt one of the three mutually exclusive identities.

**Figure 7:**
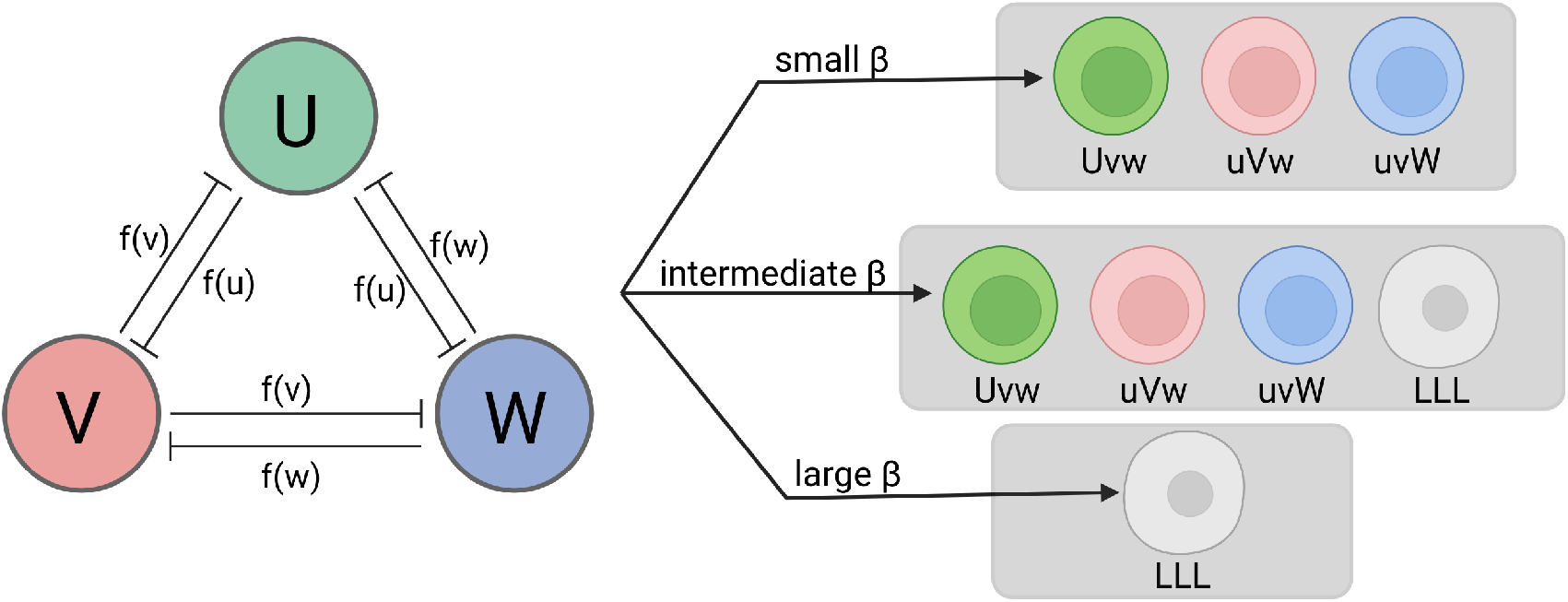
Phenotypic regimes controlled by the degradation rate *β*. *(Left)* The genetic toggle triad network motif, where master regulators *U* (green), *V* (red), and *W* (blue) engage in symmetric mutual repression. *(Right)* The landscape of accessible cellular fates dictated by *β*. **Top:** Small *β* values force commitment into one of three distinct differentiated phenotypes (*Uvw, uV w, uvW* ), corresponding to the green, red, and blue cells, respectively, where a single master regulator takes over. **Center:** Intermediate *β* values establish a multistable regime allowing for the coexistence of both the progenitor and all three differentiated lineages, providing a basis for lineage plasticity. **Bottom:** Large *β* values support only the uncommitted stem cell state (**LLL**), depicted in light grey.

Ultimately, this analysis establishes two key insights. First, it demonstrates the robustness of mutual repression as a mechanism even for three competing nodes, which reliably resolves into single-positive (**HLL**) differentiated states. Second, we note the dynamical multistability in the range *β*_1_ *< β < β*_2_ which can serve as a basis for plasticity and control of developmental processes. Furthermore, Proposition 2 proves that any hybrid states (**HHL**)—where two master regulators are simultaneously highly expressed—are unstable. This re-strengthens the concept of the purely repressive triad as a robust “winner-take-all” architecture but note however that stable hybrid phenotypes (such as Th1/Th17 cells) have indeed been observed experimentally [68, 69]. Biological evidence suggests that maintaining such mixed-identity states requires additional network connections, specifically positive autoregulation [31]. While our current purely repressive model cannot support these states, we will next explore how to incorporate self-activation in the following section.

### 2.3. Tristability in Self-Activating Toggle Switch

Thus far, our modeling and analysis covered dynamics of mutual repression within two- and three-node circuits. In these architectures, a protein product strictly represses opposing genes to block their activity, as illustrated in Figs. 1 and 5. Our minimal framework successfully captured the core dynamics of these systems by first identifying the relative degradation rate *β* as the main bifurcation parameter to systematically map the resulting cell fate decisions.

However, these models have not yet accounted for the self-interaction of individual nodes. In natural gene regulatory networks, it is highly common for a transcription factor to directly regulate its own promoter [4]. This autoregulation can be either negative or positive. While negative autoregulation is known to enhance the stability of circuit outcomes and accelerate response times [70], positive autoregulation—or self-activation acts as implementation mechanism for signal amplification, facilitating multistability and long-term hysteresis [61, 71]. Crucially, self-activation is required to generate certain biological phenomena that our purely repressive models could not capture: namely, tristability in the toggle switch [72] and the stabilization of hybrid phenotypes in the toggle triad [31].

To explore how positive feedback modifies binary decision-making, we extend the classical toggle switch by incorporating self-activation. Chemically, this architecture is modeled as a reaction-kinetic system where Gene 1 and Gene 2 produce proteins *X* and *Y*, respectively. In addition to the mutual repression sites in their regulatory control regions, each gene now possesses self-activation sites that bind its own protein product (Fig. 8a). This creates the self-activating toggle switch (SATS) motif, characterized by a continuous competition between mutual antagonism and positive autoregulation (Fig. 8b). Dynamically, the outcome of this competition depends heavily on the activation threshold. If self-activation is weak or only triggers at high concentrations, the system defaults to the classical toggle logic, resolving into one of two mutually exclusive differentiated states (*Xy* or *xY* ) (Fig. 8d). However, if self-activation is strong enough to operate at lower concentrations, it can stabilize an intermediate expression level. This overcomes the exclusive differentiation from repression, giving rise to a metastable hybrid state where both *X* and *Y* stably coexist (Fig. 8c).

**Figure 8:**
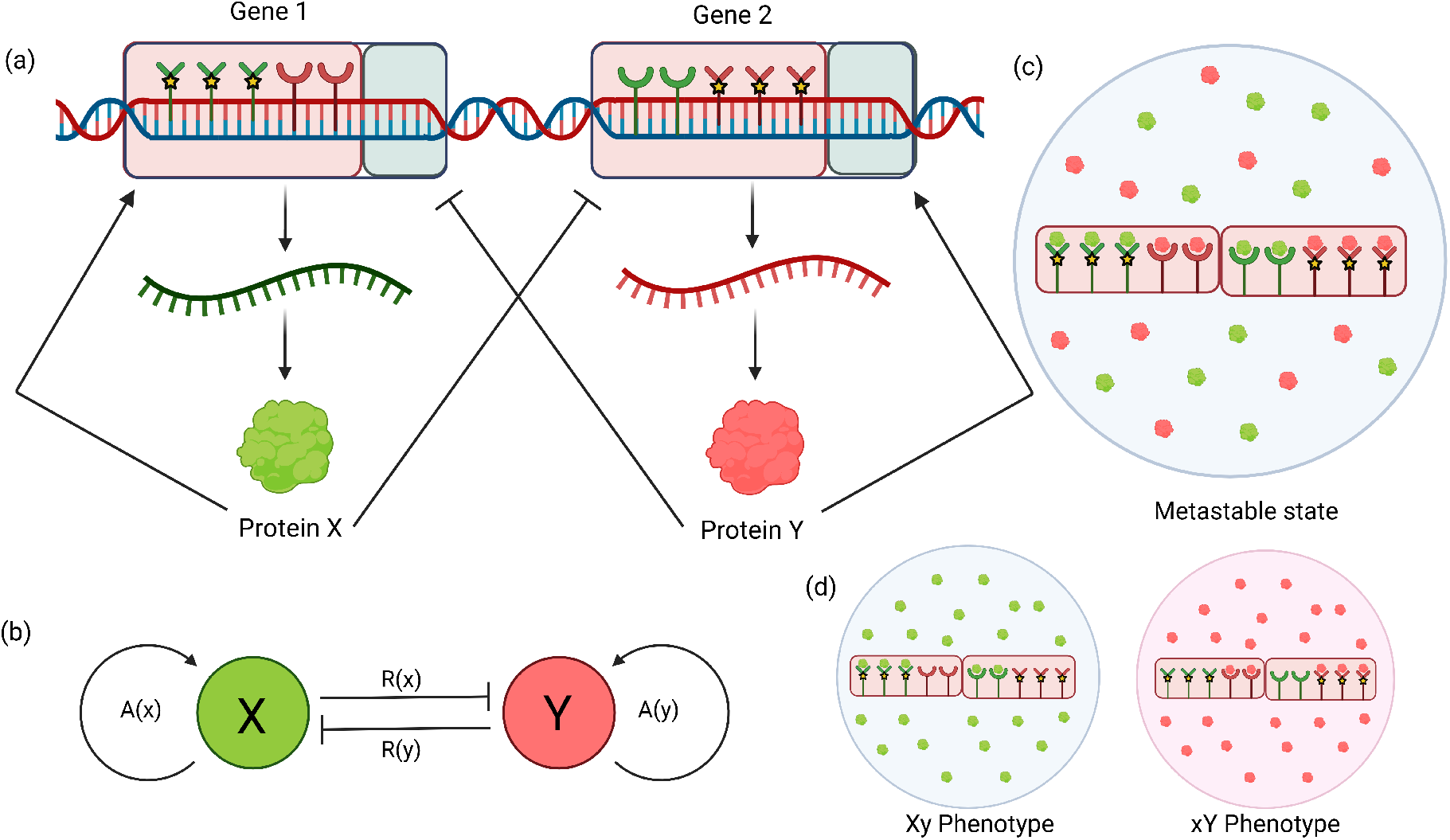
Features of Self-Activating Toggle Switch (SATS). **(a)** Each gene carries two types of regulatory binding sites: self-activation sites (starred receptors) that bind their own transcription factor and mutual repression sites (semicircle receptors) that bind the opposing transcription factor. These interactions modulate transcription, producing proteins *X* and *Y* . **(b)** Network-level abstraction of the SATS: *X* and *Y* mutually repress each other via *R*(·) and simultaneously self-activate via *A*(·). **(c)** For a low activation threshold, both genes can self-activate even at low expression levels, giving rise to a metastable hybrid state where *X* and *Y* stably coexist at intermediate levels. **(d)** For a high activation threshold, self-activation is ineffective, and the system reduces to classical toggle behavior with two mutually exclusive stable states: the *Xy* phenotype (high *X*, low *Y* ) and *xY* phenotype (low *X*, high *Y* ). Created with Biorender.com [48]

Now that we have established a picture of the gene-scale arrangement, we seek to write the governing equations for the system. The natural approach to incorporate positive autoregulation, as demonstrated by Jia *et al*. [72], is to include in the production term an additional Hill-kinetics function *A*(·) encoding self-activation. Thus to model the SATS, we add self-activation to (3) to give the resulting dynamical system:

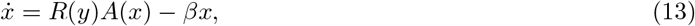

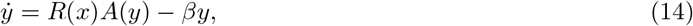

where *R*(·) is the function denoting repression by the other component while *A*(·) is the self-activation function.

For the mutual repression term, we define

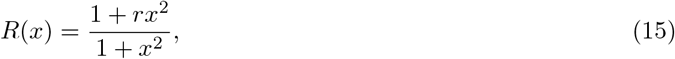

where we explicitly reintroduce the basal leakiness parameter 0 *< r <* 1. As established in theoretical models studied earlier [72, 73], this basal transcription rate is necessary for tristability. For a strongly repressed gene to eventually overcome inhibition and engage its own positive feedback loop, it must have a non-zero basal expression. The parameter *r >* 0 mathematically guarantees this survival baseline (*R*(*x*) → *r* as *x* → ∞).

To represent self-activation, *A*(·) must obey a similar Hill-like saturating form:

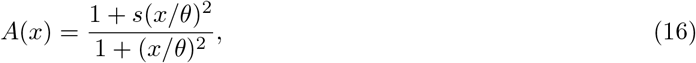

where *s >* 1 represents the maximal fold-activation strength, and *θ >* 0 is the threshold concentration for activation. At high expression (*x* ≫ *θ*), self-activation saturates to *A*(*x*) → *s*.

This form, however, leads to a parameter explosion: we must now track *r, s, β*, and *θ*. Since our goal is to specifically study the effect of self-activation, we need to identify the main tunable parameter and set reasonable values for the rest. First, we fix the basal leakiness to *r* = 0.1. Second, having already analyzed the role of *β* in detail, we fix *β* = 0.1. In a pure repressive toggle switch, this low degradation rate would guarantee bistability and ensure the uncommitted progenitor state is unstable, as we saw in Proposition 1.

We are then left to choose between *s* and *θ*, both of which control the strength of self-activation. While *s* measures the fold-increase in expression, *θ* determines the concentration threshold required for self-activation to become operational. Recalling that we normalized the mutual repression threshold (*c* = 1), *θ* is to be interpreted as the saturation threshold for self-activation compared to mutual repression. This is a useful way to conceptualize *θ*, and thus we select it as the primary bifurcation parameter for the SATS system, while setting the fold-increase in expression *s* = 20.

Consequently, tuning *θ* governs the self-activation term, and at intermediate protein concentrations:

- For a **low** activation threshold (small *θ*), *A*(*x*) → *s*.
- For a **high** activation threshold (large *θ*), *A*(*x*) → 1.

Thus, *θ* serves as the control variable which tunes the impact of positive autoregulation from dominant to absent. To visualize this parameter dependence, we compute the phase space nullclines for varying activation thresholds (*θ* ∈ {8, 10, 12} ). As illustrated in Fig. 9, lower activation thresholds (strong self-activation, *θ* = 8) stabilize the central metastable state, whereas higher thresholds (weak self-activation, *θ* = 12) force the system into toggle-switch-like binary differentiation. A critical stability transition occurs in the intermediate regime (plotted for *θ* = 10), where the phase space exhibits tristability—the simultaneous coexistence of both the asymmetric differentiated states and the symmetric hybrid state.

**Figure 9:**
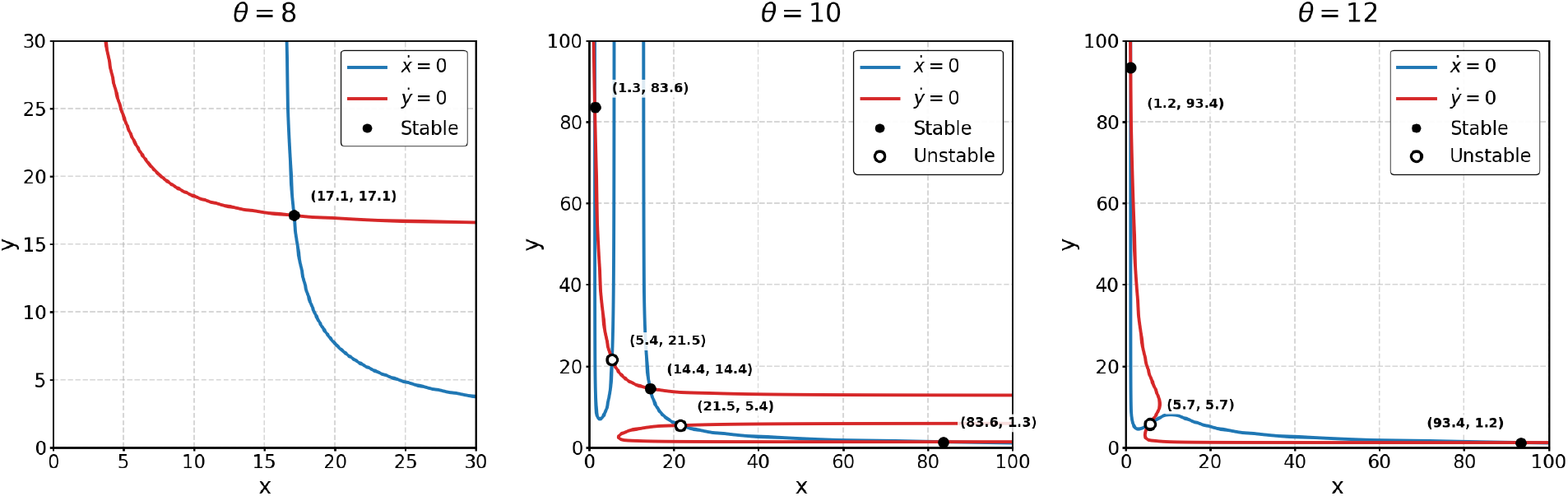
Nullclines and critical points for varying activation thresholds. (a) For *θ* = 8, strong self-activation stabilizes the central hybrid state. (b) For *θ* = 10, the system exhibits tristability, supporting both differentiated states and the hybrid state. (c) For *θ* = 12, self-activation is weak, and the system reduces to classical bistable differentiation. Solid circles indicate stable steady states; open circles indicate unstable saddle points.

#### Mathematical Mechanism of Tristability

While the tristable regime has been extensively studied through numerical simulations [72], there remains a need for a more rigorous explanation of its occurrence. To provide this, we first draw attention to a striking geometric feature of the phase space for *θ* = 10 (Fig. 9b), where the nullclines intersect at a total of five points. Considering the behavior of the *x*-nullcline from the symmetric critical point towards *x* = 0, it first reaches toward infinity, folds back to intersect at the unstable saddle point, and finally reverses direction again to intersect the *y*-nullcline at the asymmetric critical point on its final asymptotic trajectory toward infinity. Such non-monotonic behavior is highly unusual in simple regulatory models, and we term this severe geometric deformation “snaking.”

Far from being a mere pathological artifact, we establish that this snaking is precisely the geometric mechanism that permits the coexistence of multiple stable states. Without this topological deformation, the nullclines would remain far too monotonic to yield five distinct intersections. To understand this behavior in detail, we consider the *x*-nullcline:

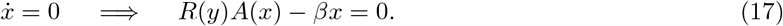

We examine the behavior of this nullcline in the asymptotic limit (*y* → ∞). In this limit of strong repression from the opposing gene, the repression function reduces to its basal leakiness, *R*(*y*) → *r*. The steady-state condition then simplifies to:

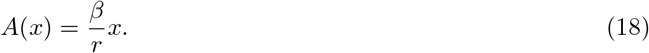

This formulation provides crucial justification for the necessity of *r >* 0; without this basal leakiness, the steady-state equation would collapse, precluding any snaking behavior, as previously studied in the literature [73].

Substituting the explicit Hill-kinetics form of the activation function from Eq. (16) yields:

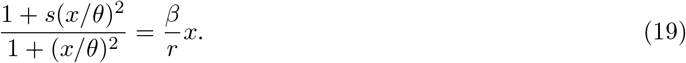

Clearing the denominator and expanding the terms gives:

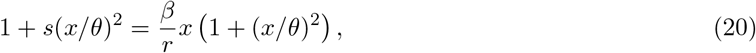

which rearranges into a standard cubic polynomial equation:

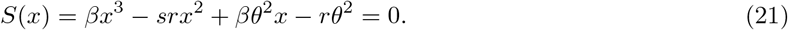

The positive real roots of Eq. (21) determine the asymptotic behavior of the nullcline at large *y*. A single root implies a unique asymptote, whereas the existence of three positive roots implies three distinct asymptotes, providing the necessary geometric condition for nullcline snaking. While snaking alone does not strictly guarantee tristability, it is the primary structural cause for the five intersection points (three stable, two unstable) required to achieve it.

This cubic equation, *S*(*x*) = 0, is plotted in Fig. 10 for the three values of *θ* considered in Fig. 9. As expected, only for the intermediate value (*θ* = 10) does *S*(*x*) admit the three roots necessary to support tristability in this circuit. Furthermore, we note that the roots of this cubic actually serve as an excellent approximation for the steady-state expression levels of the critical points in the full SATS system (13)–(14).

**Figure 10:**
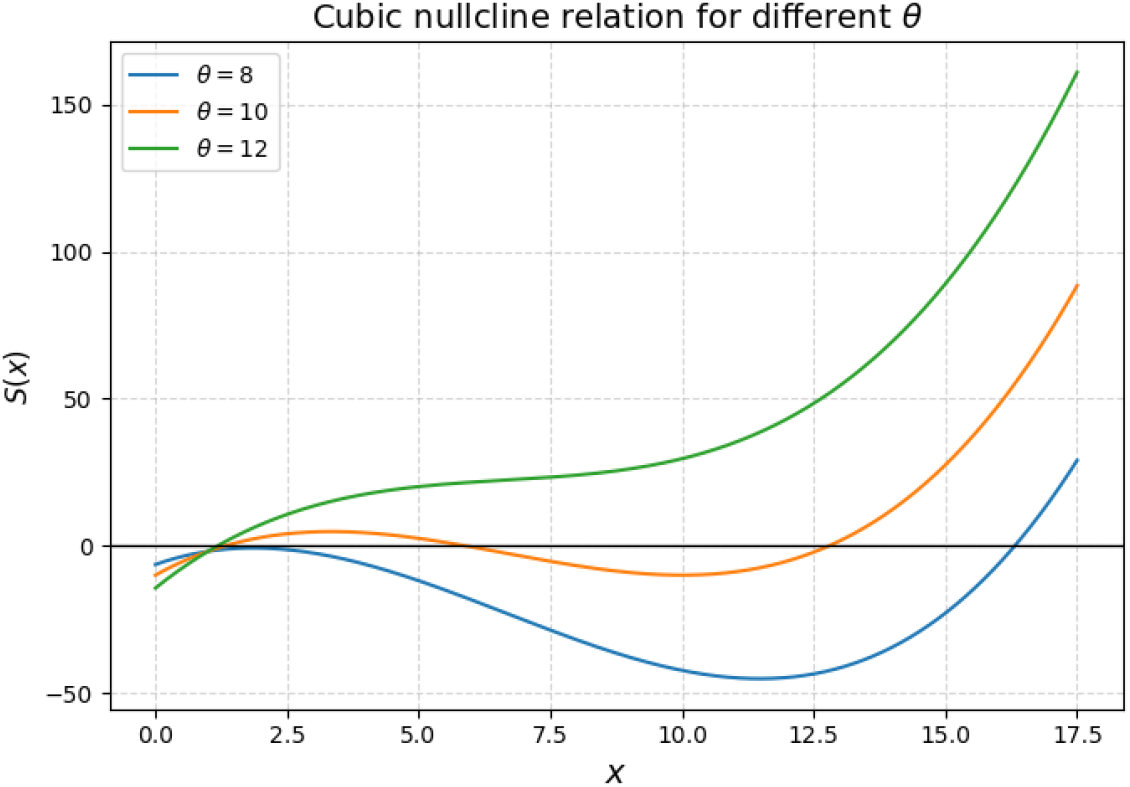
Asymptotic nullcline geometry governed by the roots of *S*(*x*). The cubic polynomial *S*(*x*) = *βx*^3^ − *srx*^2^ + *βθ*^2^*x* − *rθ*^2^ defines the asymptotic intersections of the system’s nullclines. Evaluated for *θ* ∈ {8, 10, 12}, the root multiplicity influences the number of steady states. Specifically, only the intermediate regime (*θ* = 10) admits three positive real roots, establishing a strong heuristic explanation for the asymptotic snaking of the nullclines and the emergence of tristability.

Having established this algebraic heuristic for the onset of tristability, we have demonstrated that our symmetric framework is not strictly limited to pure mutual repression, but can be successfully extended to incorporate additional mechanisms such as positive autoregulation. While there is clear potential to generalize this formalism to larger networks with highly complex node interactions—a direction we explore at the conclusion of this paper—we now shift our focus to a fundamentally different physical problem: spatiotemporal pattern formation.

## 3. From Network Motifs to Spatiotemporal Patterns

Thus far, we have established the various types of temporal multistability that can emerge from local network motifs within an isolated cell. However, functional tissue development requires these distinct phenotypes to achieve precise, stable positional arrangements [74, 75]. To investigate whether the multistability generated by these local ODE models can be robustly maintained and patterned across a continuous domain, we must transition to a spatiotemporal framework. In vivo, cells coordinate this spatial organization through continuous intercellular communication via juxtacrine contacts [76], gap junctions, or secreted morphogens [77]. To construct a minimal physical model of this process, we abstract these complex pathways into a single transport mechanism: linear spatial diffusion. While master transcription factors rarely exit the nucleus directly [33], they govern downstream cascades of secreted ligands that reliably broadcast the cell’s internal state to its neighbors [35]. Therefore, we phenomenologically model this intercellular communication as the “effective diffusion” of the master regulators themselves.

The classical mathematical framework for diffusion-driven pattern formation is the Turing instability [75]. However, Type-I Turing mechanisms strictly demand specific activator-inhibitor kinetics and significantly unequal diffusion coefficients [78]—conditions explicitly ruled out by our symmetric topology. Furthermore, as demonstrated by Roy *et al*. [79], the uniform steady states of these multistable toggle architectures are homogeneously stable against spatial perturbations.

Thus, we propose that biological pattern formation may arise from a more intuitive mechanism: a spatial “tug-of-war” [36] between competing phenotypes. Given that the intrinsic network topology establishes a discrete set of stable phenotypes, we ask a direct question: can pure spatial diffusion allow for the formation of interfaces between them? Furthermore, what are the exact stability properties of these interfaces as distinct domains compete across a tissue?

To address this, we embed our ODE networks into a bounded spatial domain Ω ⊂ ℝ^*d*^ subject to homogeneous Neumann (zero-flux) boundary conditions, mimicking an isolated *in vitro* tissue culture. The subsequent sections establish a strict topological hierarchy governing these interfaces. In **Section 3.1**, we prove that two-node networks are topologically constrained; one gene inevitably outcompetes the other across the entire domain. Consequently, any heterogeneous pattern is inherently transient, although its lifetime can be significantly extended via self-activation. Conversely, in **Section 3.2**, we demonstrate that introducing a third competitive node in the Toggle Triad lifts this restriction. This extra degree of freedom allows the spatial tug-of-war to support stable coexistence of differentiated phenotypes.

### 3.1. Spatial Instability of Two-Node Networks

We begin our spatial investigation with the diffusive Toggle Switch. While engineering autonomous patterns with this motif is a major objective [80], any spatial domains formed without external gradients [81] are strictly transient. Although they may temporarily serve as pre-patterns, their boundaries inevitably drift and homogenize into a single state.

This is due to the two-node structure of the circuit. As Gomez *et al*. [36] demonstrated, adjacent domains form unstable interfaces where the majority state continually invades the weaker one, drifting until it consumes the entire spatial domain. The mathematical foundation for this instability is the Kishimoto-Weinberger theorem [82], which applies to strictly cooperative reaction-diffusion systems under Neumann boundary conditions. Because two-node competitive networks mathematically transform into cooperative systems[82],the binary toggle switch also obeys the theorem. But rather than relying on their general proof which extends to *n*-species systems, we establish the following proposition for *n* = 2 species to show that diffusive toggle switch cannot autonomously sustain spatial patterns.

##### Proposition 3.

*Instability of Spatially Heterogeneous Equilibria*

*Consider the symmetric Toggle Switch reaction-diffusion system defined on a bounded, convex spatial domain* Ω ⊂ ℝ^*d*^ *with a smooth boundary* ∂Ω, *subject to Neumann boundary conditions:*

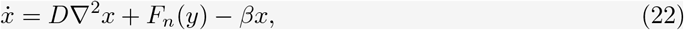

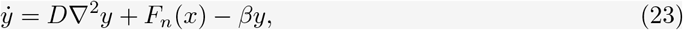

*where the repression kinetics are governed by the function F*_*n*_(*u*) = (1 + *u*^*n*^)^−1^ *for n >* 1. *Such a system admits no spatially stable heterogeneous solution and consequently, any stable equilibrium configuration must be homogeneous*.

*Proof*. Assume there exists a stationary solution (*x*^∗^(**r**), *y*^∗^(**r**)) defined over the domain **r** ∈ Ω. To analyze its linear stability, we introduce small perturbations of the form *x*(**r**, *t*) = *x*^∗^(**r**) + *e*^*λt*^*ψ*_1_(**r**) and *y*(**r**, *t*) = *y*^∗^(**r**) + *e*^*λt*^*ψ*_2_(**r**). Note that the homogeneous Neumann boundary conditions require the normal derivatives to vanish at the boundary, implying ∇*x*^∗^ · **n** = ∇*y*^∗^ · **n** = 0 and ∇*ψ*_1_ · **n** = ∇*ψ*_2_ · **n** = 0 for all **r** ∈ ∂Ω. Substituting these perturbations into the governing equations and linearizing about the steady state yields the eigenvalue problem:

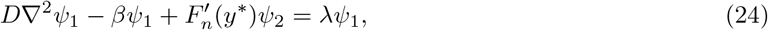

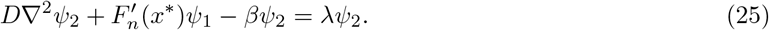

Because the function *F*_*n*_ represents repression kinetics, it is strictly monotonically decreasing, which implies 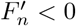 for all *u >* 0. To facilitate the stability analysis, we define the transformed perturbation vector **Ψ** = (*ψ*_1_, −*ψ*_2_)^*T*^ . This transformation allows the system to be written in the operator form ℒ**Ψ** = *λ***Ψ**,where the linear operator is defined as 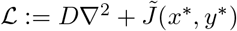. The transformed Jacobian matrix 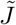 is given by:

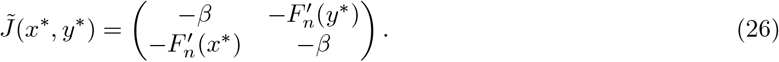

Since 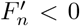, the off-diagonal terms 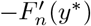 and 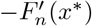 are strictly positive. Consequently, 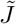 is a quasipositive matrix. This mapping makes the linearized system cooperative [82], and our next step is to consider the principal eigenvalue of this operator ℒ.

Let *X* be the Banach space of continuously differentiable vector functions on 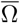, and let *K* ⊂ *X* be the solid positive cone with a non-empty interior *K*^0^. Because 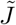 is a quasipositive matrix, we can select a sufficiently large constant *k > β* such that the spectrally shifted reaction matrix 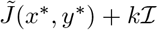 has strictly positive entries everywhere in Ω. We define the corresponding shifted differential operator as ℒ_*k*_ = ℒ + *k*ℐ . Because the diffusion operator *D*∇^2^ is uniformly elliptic and the shifted reaction matrix is fully positive, the strong maximum principle dictates that the resolvent operator (*µ*ℐ − ℒ_*k*_)^−1^ is compact and strongly positive [83]. Specifically, the resolvent maps any non-zero vector function from the cone *K* strictly into its interior *K*^0^.

By the Krein-Rutman theorem, the resolvent has a real, simple principal eigenvalue [84] with eigenvector in the cone *K*. Mapping this spectral radius back to the original differential operator formally establishes that ℒ possesses a dominant principal eigenvalue *λ*_*N*_ . Furthermore, the theorem guarantees that the corresponding principal eigenfunction under Neumann boundary conditions, **Ψ**_*N*_, is unique and strictly positive, meaning **Ψ**_*N*_ ∈ *K*^0^ everywhere in Ω.

To identify another eigenfunction of the linearized system, we examine the stationary reaction-diffusion equations satisfied by the heterogeneous solution:

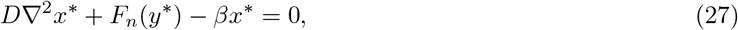

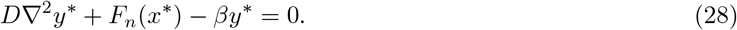

Because these equations hold everywhere in the interior of the domain Ω, we differentiate them with respect to the spatial coordinates. Applying the gradient operator ∇ to both sides yields:

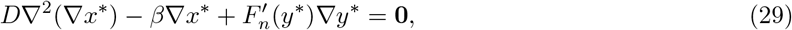

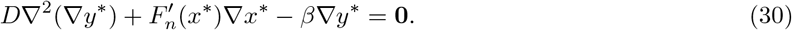

Recalling our cooperative transformation 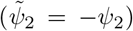, we construct the transformed spatial gradient vector ***ϕ*** = (∇*x*^∗^, −∇*y*^∗^)^*T*^ . Substituting this into our differentiated equations perfectly recovers the linearized eigenvalue problem for *λ* = 0:

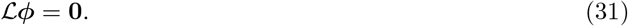

Thus, the spatial gradient ***ϕ*** is an eigenfunction associated with the zero eigenvalue. Because the original stationary solution follows homogeneous Neumann conditions (∂_*n*_*x*^∗^ = ∂_*n*_*y*^∗^ = 0) and tangential variations are zero at the interface boundary, the full spatial gradient is forced to vanish at the domain walls: ***ϕ***(**r**) = **0** for **r** ∈ ∂Ω. Consequently, this zero-eigenvalue eigenfunction ***ϕ*** strictly satisfies Dirichlet boundary conditions.

We must now determine the principal eigenvalue *λ*_*N*_ for our physical perturbations, which follow Neumann boundary conditions. The variational principle (Rayleigh-Ritz) [85] dictates that expanding the permissible function space from the restrictive Dirichlet conditions (*ψ* = **0** on ∂Ω) to the less restrictive Neumann conditions (∂_*n*_*ψ* = **0** on ∂Ω) can only increase or maintain the maximum eigenvalue. Thus, *λ*_*N*_ ≥ *λ*_*D*_ = 0.

Assume, for the sake of contradiction, that *λ*_*N*_ = *λ*_*D*_ = 0. Because the Krein-Rutman theorem guarantees that the principal eigenvalue is simple (multiplicity one), the strictly positive Dirichlet eigenfunction ***ϕ*** would also be forced to perfectly solve the Neumann eigenvalue problem. However, ***ϕ*** is strictly positive in the interior and equals zero at the boundary (its absolute minimum). Hopf’s Boundary Lemma [85] asserts that at this boundary minimum, the outward normal derivative must be strictly negative:

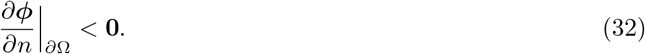

This strictly negative normal derivative explicitly violates the zero-flux requirement (∂_*n*_***ϕ*** = **0**) of a Neumann eigenfunction. Therefore, the Dirichlet eigenfunction cannot serve as the principal Neumann eigenfunction. The principal eigenvalue for the Neumann problem must therefore be strictly greater than the Dirichlet maximum : *λ*_*N*_ *>* 0. Thus, there exists a set of perturbations for any heterogeneous solution which would make it linearly unstable.

#### Remark 1

(Extension to the Self-Activating Toggle Switch)

*This spatial instability extends directly to toggle switch with self-activation, defined in Eq. (13, 14). For the Self-Activating Toggle Switch (SATS), the effective Jacobian evaluated at the stationary state* (*x*^∗^, *y*^∗^) *becomes:*

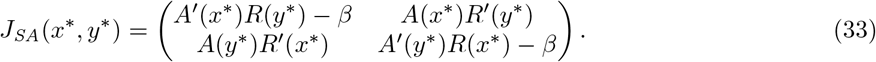

*Recall that the repressive functional form is R*(*x*) = (1 + *rx*^2^)*/*(1 + *x*^2^) *with a leakage parameter r <* 1. *Its derivative R*^′^(*x*) = 2*x*(*r* − 1)*/*(1+*x*^2^)^2^ *is strictly negative. Since A*(*x*) *>* 0, *the off-diagonal terms A*(*x*^∗^)*R*^′^(*y*^∗^) *and A*(*y*^∗^)*R*^′^(*x*^∗^) *remain strictly negative, and so applying the identical cooperative transformation* 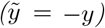 *yields again a quasipositive matrix. As before, the diagonal elements can be made strictly positive by introducing a spectral shift* ℒ_*k*_ = ℒ + *k*ℐ *for a sufficiently large constant k. Consequently, any heterogeneous solution for SATS is unstable by the exact same analysis as above*.

However, self-activation can still prolong the lifetime of these transient spatial patterns [79]. This can be understood via the variational principle [85] for the principal eigenvalue:

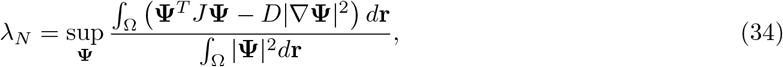

where the supremum is taken over all perturbations satisfying Neumann boundary conditions. Heuristically, the most unstable perturbation introduces asymmetry, taking the form **Ψ** ∝ (1, −1)^*T*^ . In the classical Toggle Switch (TS), which forms a sharp boundary between domains (Fig. 11a), the Jacobian *J*_TS_ evaluated at the symmetric point (the middle of the interface) yields a positive contribution, because (1, − 1)^*T*^ corresponds to an unstable eigenvector with a positive eigenvalue for the symmetric state.

**Figure 11:**
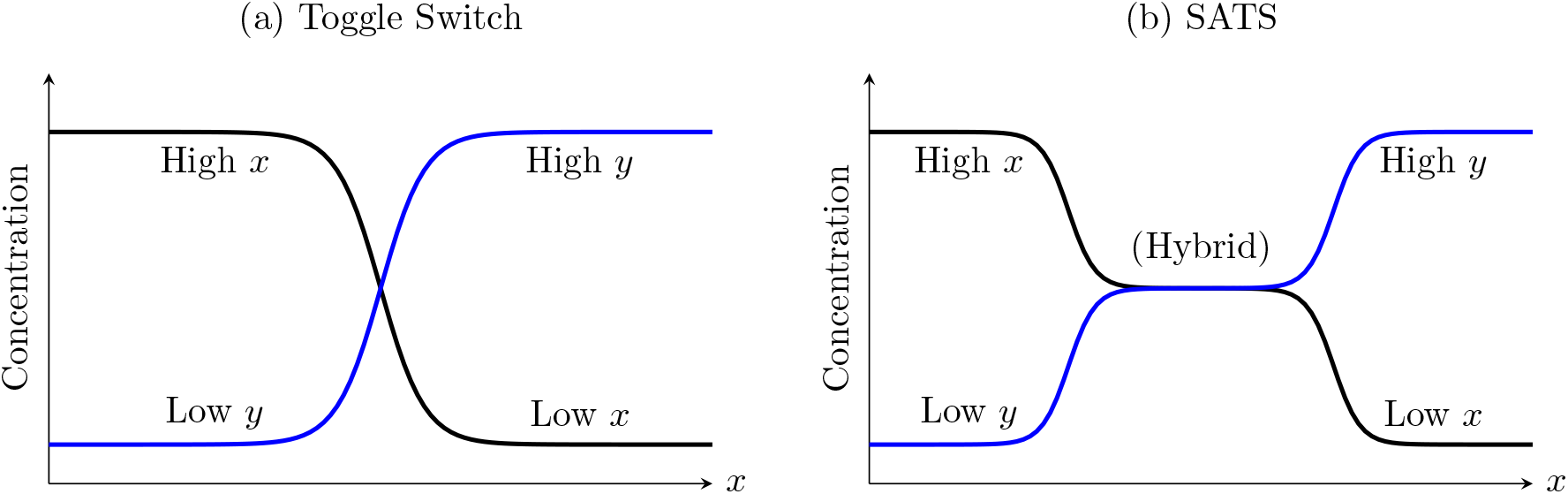
Schematic of spatial interfaces in two-node circuits. (a) The classical Toggle Switch exhibits a single, sharp interface, leading to rapid boundary migration. (b) The Self-Activating Toggle Switch (SATS) exhibits an extended spatial plateau corresponding to the stable symmetric hybrid state, which acts as a kinetic barrier that slows domain coarsening.

In contrast, for the SATS under the exact same asymmetric perturbation, the quadratic form **Ψ**^*T*^ *J*_SA_**Ψ** would be less positive due to self-activation. In fact, in the parameter regime where the SATS exhibits tristability, it may form an extended plateau of the hybrid state (Fig. 11b). Because this symmetric state is linearly stable, this quadratic term actually gives a negative contribution at the interface midpoint. Since the diffusion penalty (−*D*|∇**Ψ**|^2^) is the same for both systems for a given **Ψ**, we expect the less positive Jacobian contribution in the SATS to yield a smaller principal eigenvalue 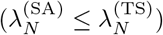, effectively making the spatial heterogeneity last significantly longer.

While mathematically unstable as *t* → ∞, these transient domains serve as vital biological pre-patterns. On finite developmental timescales, they persist long enough to trigger slower, irreversible epigenetic mechanisms that lock in tissue boundaries before the primary circuit fully coarsens [36, 86]. Phenomenologically, this boundary migration mirrors Allen-Cahn domain coarsening [38, 37]. While such phase-ordering in physical systems is generally driven by the minimization of a thermodynamic potential, here it emerges from the chemical repression of Toggle Switch. However, as we demonstrate in the next section, the addition of a third node leads to truly stable spatial patterns that escape this mathematical constraint.

### 3.2. Stable Interfaces in the Toggle Triad

Having studied patterns through two-node networks, we now investigate the three-node Toggle Triad. The addition of a third competitive node breaks the mathematical constraints of the Kishimoto-Weinberger theorem [82]. Indeed, in classical ecological models, it is well established that three-species competition-diffusion systems support stable, non-constant spatial equilibria [87, 88]. We ask whether this principle extends to gene regulatory networks to support autonomous, steady-state spatial patterning.

While spatial patterning in GRNs is less explored than classical Turing systems, prominent biological boundaries rely on mutually repressive circuits. In early *Drosophila* embryogenesis, gap genes such as *Krüppel, Giant, Knirps*, and *Tailless* [89, 40] establish sharp, non-overlapping expression boundaries along the anterior-posterior axis [42, 41, 90]. Similarly, in the vertebrate ventral neural tube, a mutually repressive triad composed of the transcription factors Pax6, Olig2, and Nkx2.2 dynamically interprets the morphogen gradient [91] to partition the tissue that ultimately generate specialized neuronal subtypes [67].

To explore the physical basis of these stable boundaries, we extend our three-node motif into a spatial domain. For analytical tractability regarding interface formation, we restrict our focus to the one-dimensional patterns, Ω = ℝ:

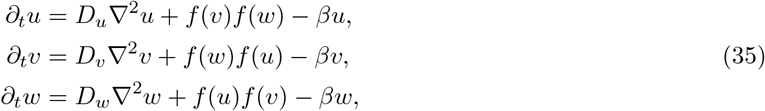

where *f* (*x*) = (1 + *x*^2^)^−1^ defines the repressive interactions.

Operating in the multistable regime (*β <* 1*/*4) where only the asymmetric states are ODE stable, numerical simulations confirm that random initial conditions rapidly resolve into stable, alternating domains of differentiated states separated by sharp interfaces (Fig. 12).

**Figure 12:**
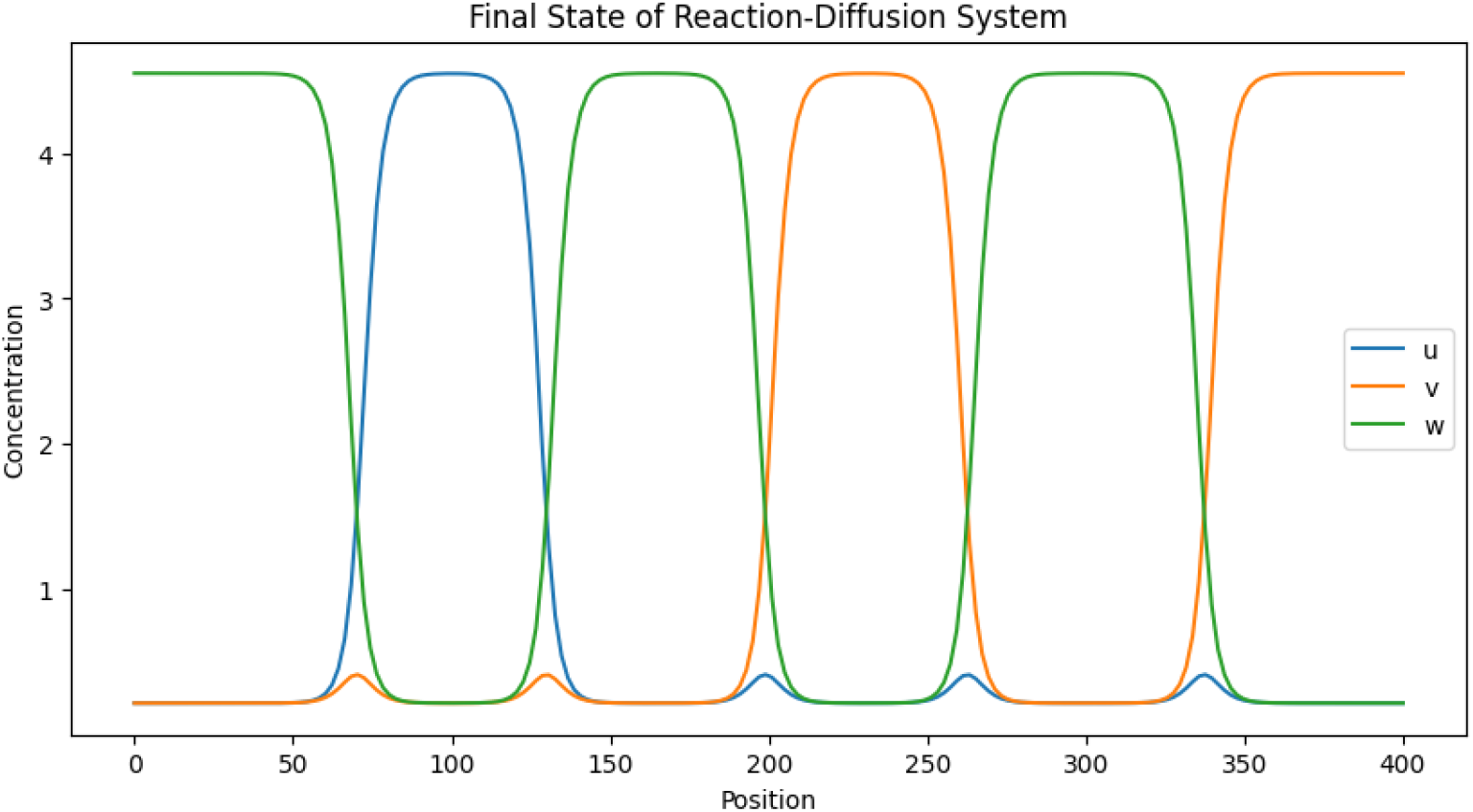
One-dimensional spatial patterning for Eq. (35). Random initial conditions autonomously evolve into stable domains of distinct phenotypic states, separated by stable interfaces (*D*_*u*_ = *D*_*v*_ = *D*_*w*_ = 1).

Mathematically, these stationary interfaces correspond to heteroclinic orbits connecting distinct homogeneous equilibria. Specifically, an interface between a high-*u* and high-*v* domain corresponds to a spatial trajectory (*u*(*x*), *v*(*x*), *w*(*x*)) satisfying the asymptotic boundary conditions:

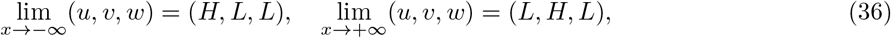

along with vanishing spatial derivatives at infinity (∂_*x*_*u* = ∂_*x*_*v* = ∂_*x*_*w* = 0 as *x* → ±∞). Numerical integration explicitly confirms the existence of these heteroclinic orbits. Therefore, we focus our analytical efforts on extracting the structural properties of the interface midpoint.

To visualize this structure, we schematically depict the spatial profile in Figure 13. The dominant species *u* and *v* exchange concentrations monotonically across the interface, intersecting symmetrically at a midpoint (*ϕ, ϕ*). Crucially, the third species *w* remains basally repressed across the entire bulk domain, exhibiting only a localized spatial bump of maximum height *c*_*m*_ near the interface core.

**Figure 13:**
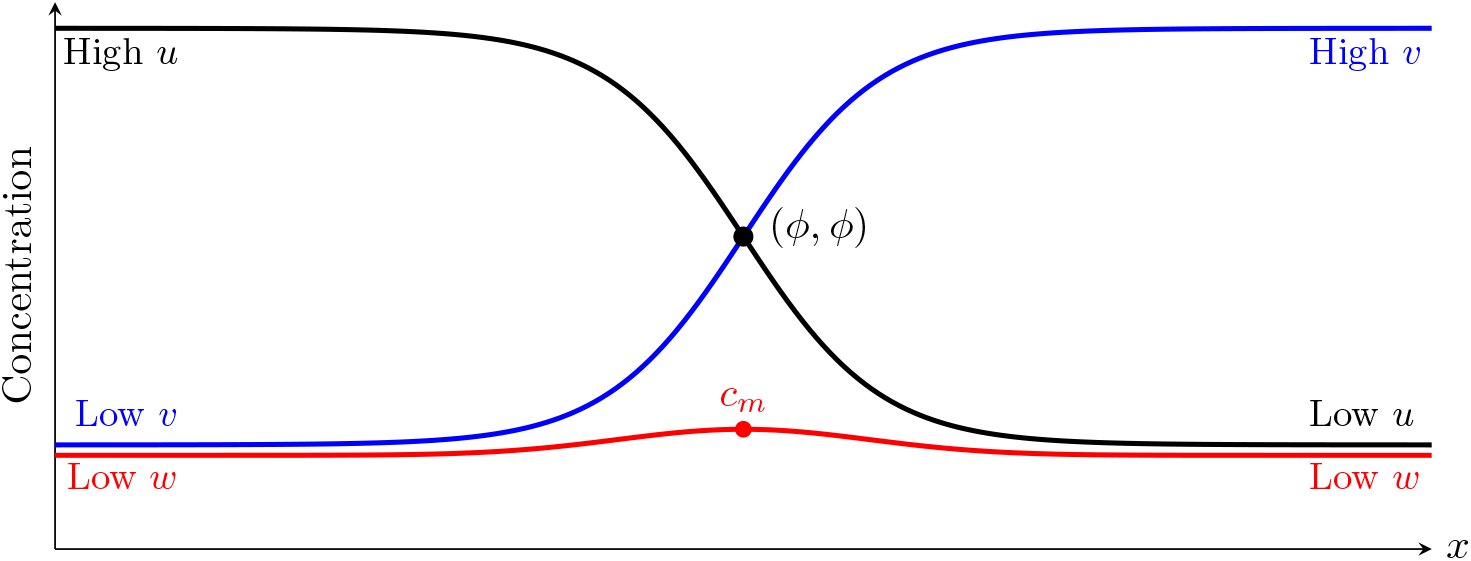
Schematic interface between high-*u* and high-*v* domains. Species *u* (black) decreases and *v* (blue) increases across the spatial interface, intersecting symmetrically at (*ϕ, ϕ*). The third species *w* (red) remains repressed in both bulk domains but exhibits a characteristic spatial bump of maximum height *c*_*m*_ at the interface center.

By repeating and concatenating this profile across space, alternating domains of high-*u* and high-*v* states emerge as the fundamental building blocks of the striped patterns observed numerically. Because *w*(*x*) remains near its low equilibrium state across interface of other two genes, its spatial gradient in the heteroclinic interface in Fig. 13 is negligible. We can therefore mathematically isolate the interface dynamics between the two active competitors by taking the singular limit *D*_*w*_ → 0, while maintaining *D*_*u*_ = *D*_*v*_ = *D* to preserve the fundamental spatial symmetry of the competing states.

##### Proposition 4

*Let β < β*_1_ *and suppose D*_*u*_ = *D*_*v*_ = *D, D*_*w*_ = 0. *If the toggle triad PDE system* (35) *admits a heteroclinic interface solution satisfying the boundary conditions* (36), *then at the midpoint x* = 0 *the solution must satisfy*

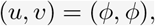

*where ϕ is a root of*

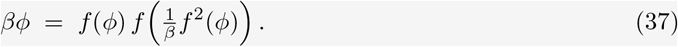

*The homogeneous state* ***LLL*** *which satisfies f*^2^(*L*) = *βL satisfies* (37) *trivially*.

*Proof*. Consider the toggle triad PDE system (35) with *D*_*u*_ = *D*_*v*_ = 1, *D*_*w*_ = 0:

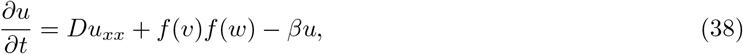

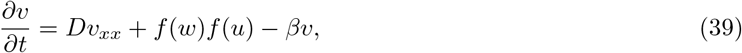

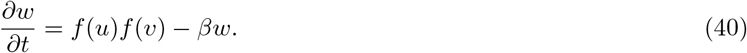

For stationary solutions:

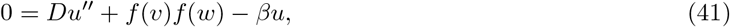

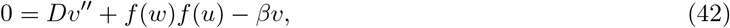

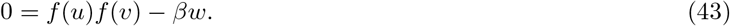

Eliminating *w*(*x*) using the third equation gives

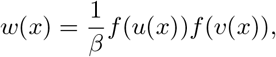

and substituting this into the first two equations yields a reduced two-variable system:

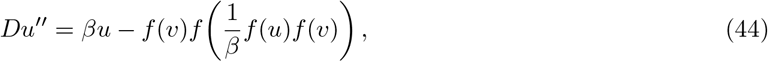

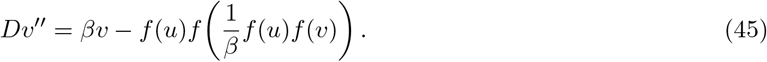

This system is symmetric under (*u*(*x*), *v*(*x*)) ↦ (*v*(−*x*), *u*(−*x*)), so the interface midpoint *x* = 0 satisfies

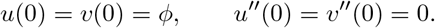

Substituting these midpoint conditions gives the algebraic relation

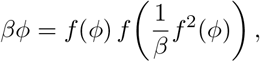

which is exactly (37).

The homogeneous state **LLL** (*ϕ* = *L*) satisfies *βL* = *f* ^2^(*L*) and therefore is a trivial solution of (37).

#### Remark 2

*Equation* (37) *has three numerical roots. Of these, the physically realized interface midpoint corresponds to ϕ* = *L (the LLL value). The two other roots are spurious and do not correspond to any realizable interface state*.

## 4. Conclusion

We conclude our systematic study of three simple regulatory motifs. In an era dominated by large-scale gene network simulations [31, 92], the utility of minimal pen-and-paper models might seem questionable, but the explosion of computational complexity actually demands systematic abstraction. Idealized models, grounded in clear physical assumptions and analytical tractability, provide the basic intuition required to understand actual *in vivo* developmental dynamics. Our findings, summarized in Table 1, map the behavior of these motifs across various parameter regimes and connect their temporal multistability to spatiotemporal organization.

**Table 1:**
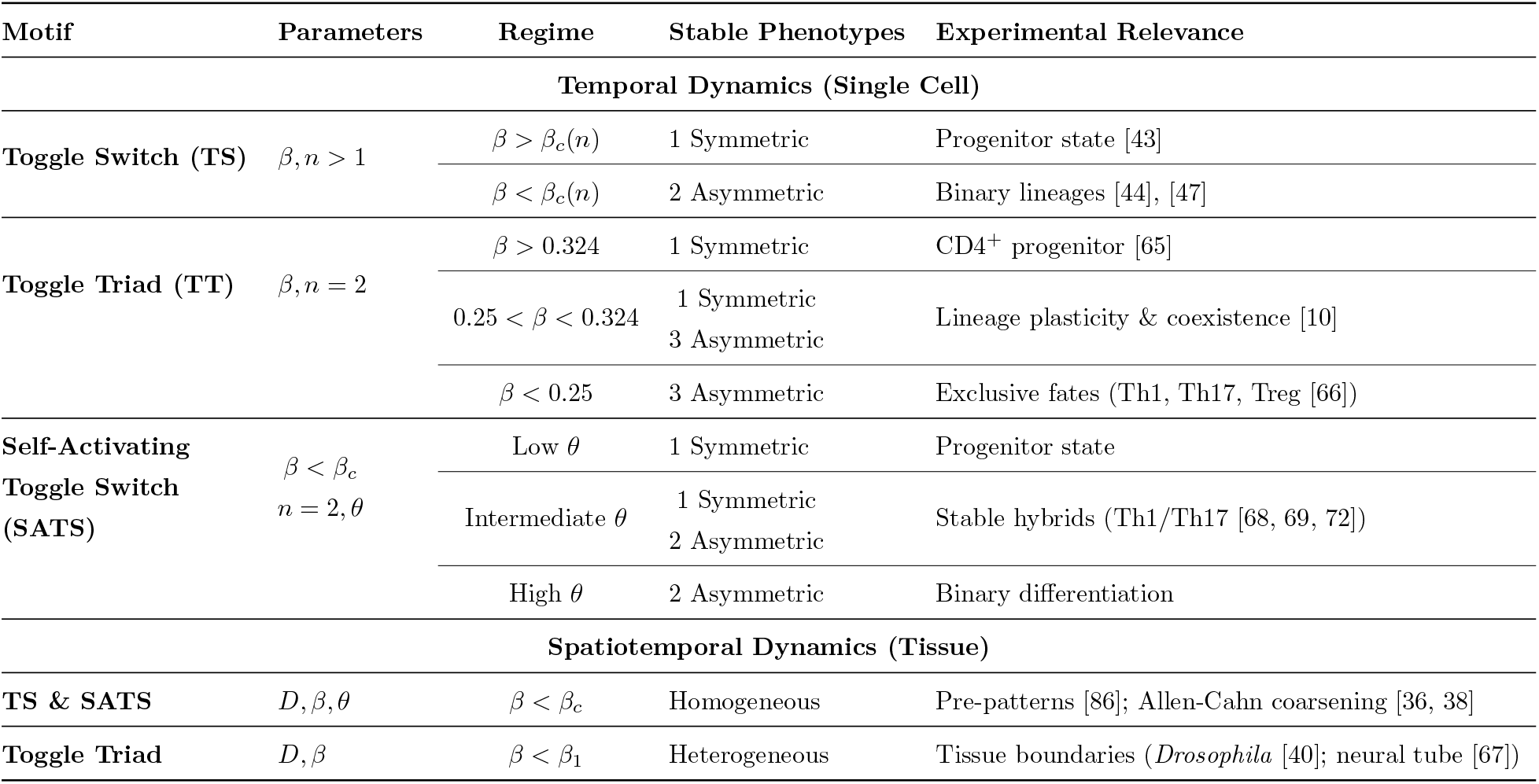
Summary of Network Motifs and Dynamical Behaviors. The exact analytical and biological outcomes of the minimal regulatory networks.

### Universal Control of Cell Fate

Moving beyond the developmental “statics” captured by parameter-randomized ensembles [31], our framework explicitly tracks both the symmetric progenitor and asymmetric mature states to map the bifurcations that drive lineage commitment. Through this approach, we first isolated the relative degradation rate, *β*, as the universal control parameter for this transition within purely repressive networks. High protein degradation prevents sufficient repressor accumulation, locking cells in an inactive, uniform progenitor state. Differentiation is triggered exclusively when *β* drops below a critical threshold. This analytical requirement perfectly aligns with recent experimental evidence showing that global protein degradation acts as a biochemical clock, dictating the species-specific “tempo” of embryogenesis [93]. Just as tuning bulk protein turnover physically scales developmental timelines *in vivo*, crossing this theoretical *β* threshold serves as the fundamental mathematical trigger for exiting pluripotency.

Building directly upon this *β* baseline, adding positive autoregulation allowed us to systematically isolate the activation threshold, *θ*, as the next primary control parameter. In the Self-Activating Toggle Switch (SATS), *θ* drives the emergence of tristability. This can be geometrically explained by analyzing the asymptotic behavior of the nullclines, rigorously demonstrating why only intermediate values of *θ* can support hybrid states alongside differentiated ones. We can extend this logic to the three-node Toggle Triad to argue that self-activating nodes support hybrid double-positive states [31]—such as a co-expressed *UV w* phenotype acting as a metastable bridge between the committed *Uvw* and *uV w* attractors. If node *w* remains basally repressed, the active nodes (*u* and *v*) effectively decouple into an isolated SATS subsystem.

### Spatiotemporal Self-Organization

Extending these multistable phenotypes into a spatiotemporal domain argues that robust developmental patterning often arises not from classical Turing instabilities [78], which require restrictive differential diffusion rates, but from the spatial arrangement of competing phenotypes [36]. Coupling multistable reaction kinetics with simple spatial diffusion establishes a reaction-diffusion system that drives pattern formation through heteroclinic interfaces.

As demonstrated by our exact derivations, these spatially extended networks present fundamental topological constraints. Two-node interfaces are strictly transient, and true spatial stability requires the topological freedom of a third competitive node. The Toggle Triad lifts this mathematical constraint, presenting deeply challenging mathematical problems where established analytical tools for heteroclinic orbits [94] will prove invaluable. Moving forward, translating these theoretical insights from idealized toy models into functional circuits in synthetic biology labs will require devising increasingly clever mathematical and experimental techniques.

### From Minimal Motifs to Complex Regulatory Landscapes

Looking forward, our exact analytical framework provides a robust foundation for exploring higher-order cellular decision-making. Natural topological extensions include coupling toggle switches into circular polygons and polyhedral motifs, such as the toggle tetrahedron (Fig. 14a-c). Analyzing these symmetric networks with both positive and negative autoregulation could rigorously explain the unexpected multi-positive configurations observed in recent numerical studies [95, 92].

**Figure 14:**
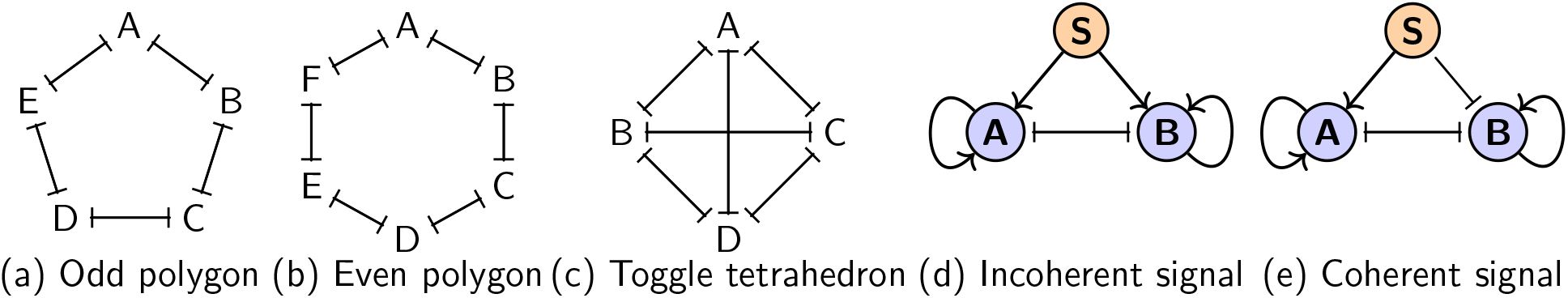
Representative motifs extending the toggle framework. Higher-order network topologies such as (a) odd polygons, (b) even polygons, and (c) the toggle tetrahedron exhibit rich, multi-positive steady states. Additionally, external regulatory nodes (**S**) can disrupt internal symmetry via (d) incoherent signaling, which stabilizes hybrid states, or (e) coherent signaling, which actively forces lineage commitment.

Furthermore, *in vivo* decision-making networks are constantly subjected to external signals that break structural symmetry (Fig. 14d-e). Integrating external bias parameters [72] into our models will allow us to understand how local tissue environments coherently force asymmetric lineage commitment [96] or incoherently stabilize uncommitted phenotypes [30]. While assuming perfect node symmetry was mathematically essential to isolate the uncommitted state, *in vivo* networks possess inherent parametric asymmetries. Developing analytical tools to perturb this symmetry and study how biochemical biases influence decision-making is a critical next step. Furthermore, our spatiotemporal models must be extended to incorporate detailed communication mechanisms—such as chemotaxis [97], anisotropic diffusion [98], and mechanochemical coupling [99]. Alongside numerical simulations in higher dimensional domains, varying initial conditions might explain how multistable networks effectively discretize continuous morphogen gradients into sharp spatial boundaries [91].

Gene regulatory networks have evolved alongside life through billions of years, yet the physical principles involved—spontaneous symmetry breaking, repressive and activating interactions, and pattern formation—are remarkably simple and universal. By understanding these basic operating principles, modern research in systems and synthetic biology continues to uncover exactly how life develops its endlessly beautiful forms.

## Acknowledgments

UR acknowledges the support of DST-INSPIRE Faculty Fellowship (DST/INSPIRE/04/2022/003052), Government of India. MKJ acknowledges the support of the Param Hansa Philanthropies. TB and AK would like to thank Thomas Hillen for mentorship and many insightful discussions.

## A. Proof of Proposition 1

*Proof*. To analyze local stability, we compute the Jacobian of system (3):

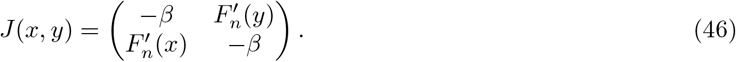

Evaluating at the symmetric equilibrium (*ϕ, ϕ*) gives

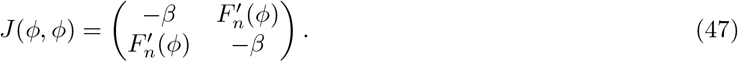

The characteristic equation of *J*(*ϕ, ϕ*) is

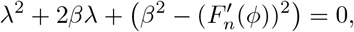

with eigenvalues

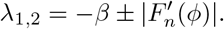

A change in stability occurs when *λ* = 0, i.e.

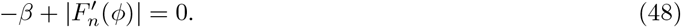

Since 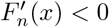 for *x >* 0, we have 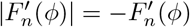, so condition (48) becomes

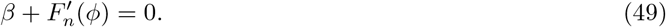

Using the equilibrium condition, we substitute 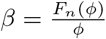 into (49). Since

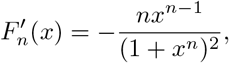

we obtain

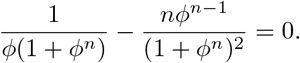

Multiplying by *ϕ*(1 + *ϕ*^*n*^)^2^ yields

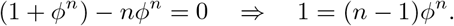

Thus,

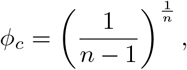

and substituting into 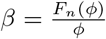 gives

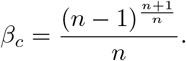

Because the destabilizing eigenvector is antisymmetric, this loss of stability results in a pitchfork bifurcation, creating two asymmetric equilibria.

## B. Proof of Proposition 2

*Proof*. **1. Symmetric State** (*ϕ, ϕ, ϕ*). At a fully symmetric fixed point, all nodes are equal: *u* = *v* = *w* = *ϕ*. The equilibrium condition is given by (10). Let us define:

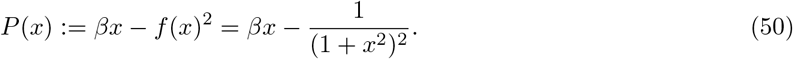

At *x* = 0, *P* (0) = −1; as *x* → ∞, *P* (*x*) → + ∞ . Since *βx* is strictly increasing and *f* (*x*)^2^ is strictly decreasing for *x >* 0, *P* (*x*) is strictly monotonically increasing. Thus, there exists exactly one positive root *ϕ*.

To determine stability, we evaluate the Jacobian at (*ϕ, ϕ, ϕ*):

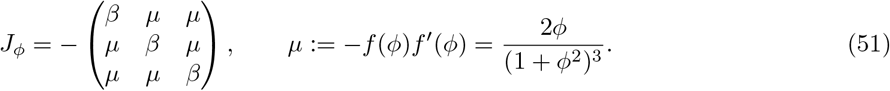

Its eigenvalues are:

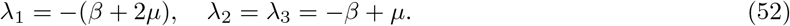

Since *µ >* 0, the first eigenvalue is always negative. Stability requires *λ*_2_ = *λ*_3_ *<* 0, yielding:

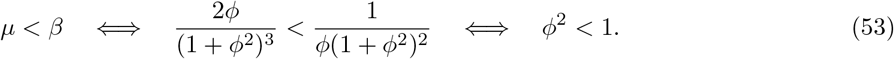

As *β* decreases, the intersection of *βx* and *f* (*x*)^2^ shifts to larger values of *ϕ*. The critical point *ϕ* = 1 occurs exactly at *β* = 1*/*4. Therefore, the symmetric low-expression LLL state is linearly stable if and only if:

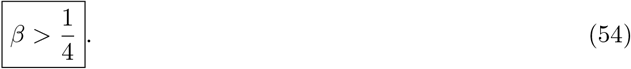

### 2. Asymmetric States

Let (*u*^∗^, *v*^∗^, *w*^∗^) satisfy the equilibrium conditions:

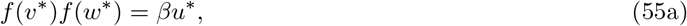

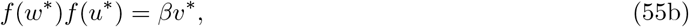

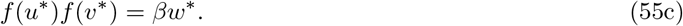

Dividing each equation by its respective right-hand variable and multiplying by *f* (*u*^∗^), *f* (*v*^∗^), and *f* (*w*^∗^) yields a common constraint:

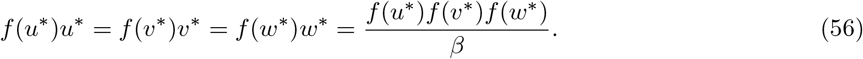

We define a constant *η* for this shared value:

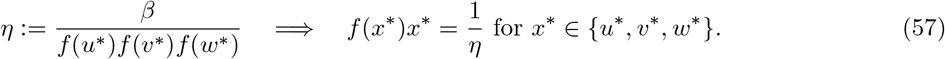

Substituting 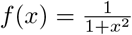, this simplifies to:

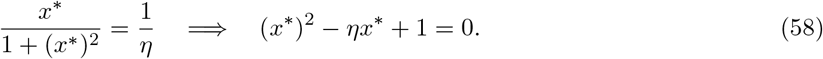

This quadratic yields two positive roots if and only if *η* ≥ 2:

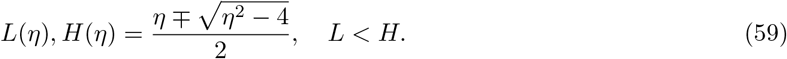

By Vieta’s formulas, these roots satisfy the crucial symmetric properties *HL* = 1 and *H* + *L* = *η*.

#### Stability of (*H, L, L*)

Evaluating the Jacobian at (*H, L, L*) reveals a remarkable symmetry. Because *HL* = 1, we have *f* ^′^(*H*)*f* (*L*) = *f* ^′^(*L*)*f* (*H*). This allows us to define the Jacobian as:

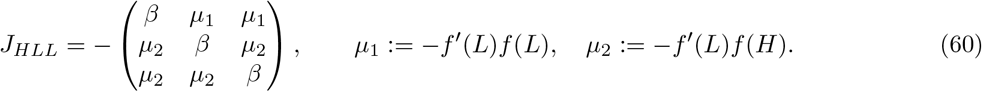

Using *f* (*L*)*L* = 1*/η, H* = 1*/L*, and the equilibrium conditions, we can rewrite the parameters strictly in terms of *H* and *η*:

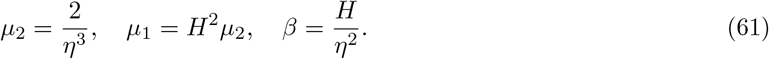

The eigenvalues of *J*_*HLL*_ are:

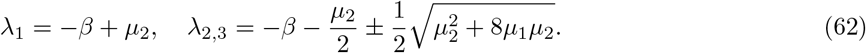

Stability is governed by the largest root, *λ*_3_. Substituting our *H* and *η* expressions into *λ*_3_ *<* 0 gives:

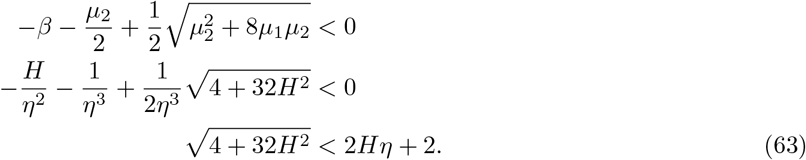

Substituting 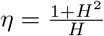, the right-hand side becomes 4 + 2*H*^2^. Squaring both sides and simplifying yields:

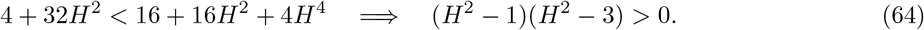

Since we are on the high branch (*H >* 1), this requires 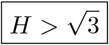.

Using 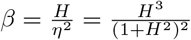, we find the critical bifurcation parameter at 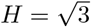:

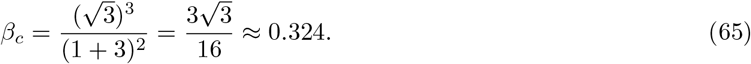

Therefore, the asymmetric state (*H, L, L*) is linearly stable if and only if:

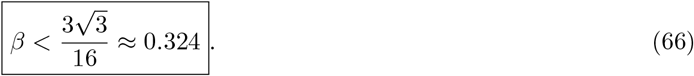

### 3. Instability of (*H, H, L*). To verify that (*H, H, L*) is never stable, we define

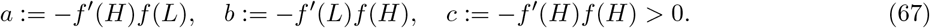

The Jacobian evaluated at (*H, H, L*) is:

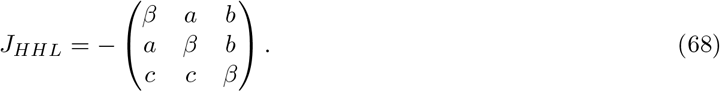

The vector (1, −1, 0)^⊤^ is an eigenvector of *J*_*HHL*_ with eigenvalue *a* − *β*. Using the steady-state relations *βH* = *f* (*H*)*f* (*L*) and *βL* = *f* (*H*)^2^, we evaluate this eigenvalue:

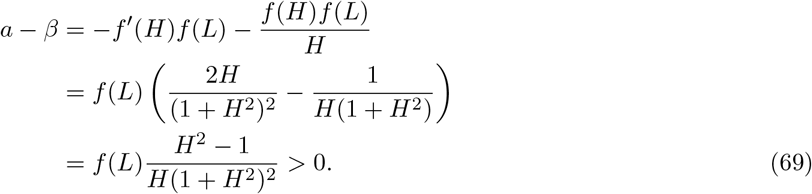

Since *HL* = 1, we have *H >* 1 which makes this eigenvalue strictly positive, proving that the (*H, H, L*) hybrid state is always unstable.

